# Pathogenic Mutations in the Kinesin-3 Motor KIF1A Diminish Force Generation and Movement Through Allosteric Mechanisms

**DOI:** 10.1101/2020.09.03.281576

**Authors:** Breane G. Budaitis, Shashank Jariwala, Lu Rao, David Sept, Kristen J. Verhey, Arne Gennerich

## Abstract

The kinesin-3 motor KIF1A functions in neurons where its fast and superprocessive motility is thought to be critical for long-distance transport. However, little is known about the force-generating properties of kinesin-3 motors. Using optical tweezers, we demonstrate that KIF1A and its *C. elegans* homolog UNC-104 undergo force-dependent detachments at ~3 pN and then rapidly reattach to the microtubule to resume motion, resulting in a sawtooth pattern of clustered force generation events that is unique among the kinesin superfamily. Whereas UNC-104 motors stall before detaching, KIF1A motors do not. To examine the mechanism of KIF1A force generation, we introduced mutations linked to human neurodevelopmental disorders, V8M and Y89D, based on their location in structural elements required for force generation in kinesin-1. Molecular dynamics simulations predict that the V8M and Y89D mutations impair docking of the N-terminal (β9) or C-terminal (β10) portions of the neck linker, respectively, to the KIF1A motor domain. Indeed, both mutations dramatically impair force generation of KIF1A but not the motor’s ability to rapidly reattach to the microtubule track. Homodimeric and heterodimeric mutant motors also display decreased velocities, run lengths, and landing rates and homodimeric Y89D motors exhibit a higher frequency of non-productive, diffusive events along the microtubule. In cells, cargo transport by the mutant motors is delayed. Our work demonstrates the importance of the neck linker in the force generation of kinesin-3 motors and advances our understanding of how mutations in the kinesin motor domain can manifest in disease.

## INTRODUCTION

The cytoskeleton of eukaryotic cells forms the structural framework for fundamental cellular processes including cell division, cell motility, intracellular trafficking, and cilia function. In most of these processes, the functional output of the microtubule (MT) cytoskeleton depends on a family of molecular motor proteins called kinesins. Kinesins are defined by the presence of a globular motor domain that contains sequences for binding ATP and MTs. Kinesins involved in intracellular trafficking use the energy of ATP hydrolysis for processive motility and force generation along the MT surface.

The kinesin-3 family is one of the largest among the kinesin superfamily and its members are primarily involved in the anterograde transport of cargoes toward the plus ends of MTs in the periphery of the cell [reviewed in (1-3)]. Genetic and microscopy studies have implicated the kinesin-3 motor KIF1A, and its orthologs, in the transport of synaptic vesicle precursors (SVPs) and dense core vesicles (DCVs) to the axon terminal (4-9). A number of inherited variants and *de novo* mutations have been identified in human *KIF1A* from clinical studies. These mutations have been linked to neurodevelopmental and neurodegenerative disorders including spastic paraplegias, encephalopathies, intellectual disability, autism, and sensory neuropathies (3, 10-15). For KIF1A-associated neurological disorder (KAND), the mutations span the entirety of the KIF1A protein sequence; the majority are located within the kinesin motor domain (aa 1-369) and are thus predicted to affect the motor’s motility properties whereas mutations located outside the motor domain are likely involved in mediating cargo binding, dimerization, and/or autoinhibition [reviewed by (3)].

Recent studies have shown that kinesin-3 proteins have striking motility properties as they are exceptionally fast and superprocessive and have dramatically higher MT-landing rates (ability to productively engage with MTs) than other kinesin motors (16-18). However, little is known about the ability of kinesin-3 motors to generate and sustain force. A general understanding of how kinesin motors generate force is largely based on studies of kinesin-1 (19-24), the founding member of the kinesin superfamily. Force generation requires the neck linker (NL), a flexible structural element that immediately follows the kinesin motor domain, which docks along the surface of the motor domain in response to ATP binding (25-28). NL docking in kinesin-1 occurs in two steps. First is the “zippering” step in which the first half of the NL (β9) interacts with β0 [the cover strand (CS)] of the core motor domain to generate a short β-sheet termed the cover-neck bundle (CNB) (29). Although formation of the CNB has been observed in structures of motor domains from kinesin-3, kinesin-5, and kinesin-6 members (30-35), its mechanical role in force generation has only been tested in kinesin-1 motors (27, 36). Second is the “latching” step where the second half of the NL (β10) interacts with surface residues of α1-β3 and β7 of the core motor domain and is latched in place via a conserved asparagine residue (the N-latch) (27, 29, 36). A role for NL latching in force generation was recently demonstrated for kinesin-1 (36). Crystal structures of kinesin-3 motor domains suggest that close contact between α1-β3 and the NL may play a role in force generation for this family as well (30, 37, 38).

Despite these structural similarities, several studies have suggested that the force-generating properties of kinesin-3 motors may be different than that of other kinesin motors. First, when forced to compete with kinesin-1, KIF1A gives up easily, suggesting that it has a high load-dependent off-rate from the MT (39-41). Second, the *C. elegans* homolog UNC-104 displays a rapid decrease in velocity and increase in MT-dissociation rate under load applied in an optical tweezers assays (42). Here, we determine the force-generating properties of two members of the kinesin-3 family, the mammalian KIF1A motor and its homolog UNC-104, present in mammalian cell lysates and purified from *E. coli* bacteria. In a single-molecule optical tweezers assay, we find that UNC-104 motors stall and then undergo detachment at an average force of 3 pN. KIF1A motors also detach at an average force of 2.7 pN but readily detach from the MT track rather than stall. Strikingly, both UNC-104 and KIF1A motors quickly reattach to the MT and resume force generation, leading to a characteristic saw-tooth force-generation pattern that is distinct from other kinesin motors to date.

To determine whether NL docking plays a critical role in force generation by KIF1A, we introduced disease-associated mutations based on their a) location in structural elements predicted to be critical for NL docking and b) mild disease phenotypes that suggest an impairment rather than loss of KIF1A protein activity. V8M and Y89D are *de novo* mutations that manifest in an autosomal dominant manner to cause pure hereditary spastic paraplegia with childhood onset [OMIM #610357, (43)] and mental retardation, autosomal dominant 9 [OMIM #614255, (44)], respectively. The V8M mutation is located in β1, immediately following the CS, and may therefore prevent CNB formation. Notably, a valine in this position is highly conserved across the kinesin superfamily [Fig S1, (45)]. The Y89D mutation is located at the α1-β3 intersection and an aromatic residue (tyrosine or phenylalanine) at this position is highly conserved across the kinesin superfamily [Fig S1, (36, 45)]. To provide insights into the molecular effects of these mutations, we performed molecular dynamics (MD) simulations, which predicted attenuating effects of both mutations on the motility and force generation of KIF1A. In optical tweezers assays, both mutations resulted in a significant decrease in force output but had no effect on the motor’s ability to rapidly reengage with the MT track. In single-molecule fluorescence assays, both mutations resulted in a decrease in speed, processivity, and landing rate on MTs under unloaded conditions. In addition, KIF1A motors containing the Y89D mutation displayed an increase in diffusive events. Finally, we used a peroxisome-targeting assay to probe the ability of WT and mutant motors to work in teams to drive organelle transport in cells. We found that mutant motors show a significant delay in organelle transport. Collectively, our results support the proposed role for the NL as a mechanical element important for kinesin motors to transport against load. Our results also provide insight into how KAND-associated mutations affect KIF1A transport in cells.

## RESULTS

### KIF1A and UNC-104 motors exhibit rapid force-dependent detachments and reattachments to MTs

To examine the force output of kinesin-3 motors, we used optical tweezers with nanometer-level spatial resolution (46, 47) to probe the force response of rat KIF1A and *C. elegans* UNC-104. As a control, we also performed experiments on the widely-studied rat kinesin-1 KIF5C. Biotinylated KIF5C(1-560)-AviTag™ motors in COS-7 cell lysates (KIF5CC) bound to streptavidin-coated trapping beads displayed typical force-generating events but frequently detached under load before reaching a stall plateau. Stalling of KIF5CC motors occurred at an average force of ~5 pN (motor stalling for ≥200 ms) (Fig 1A&C), however, the average force at which KIF5CC detaches from the MT under load is smaller (4.4 pN; Fig 1D, Table 1), consistent with previous studies (19, 24, 27, 36).

**Fig 1.**
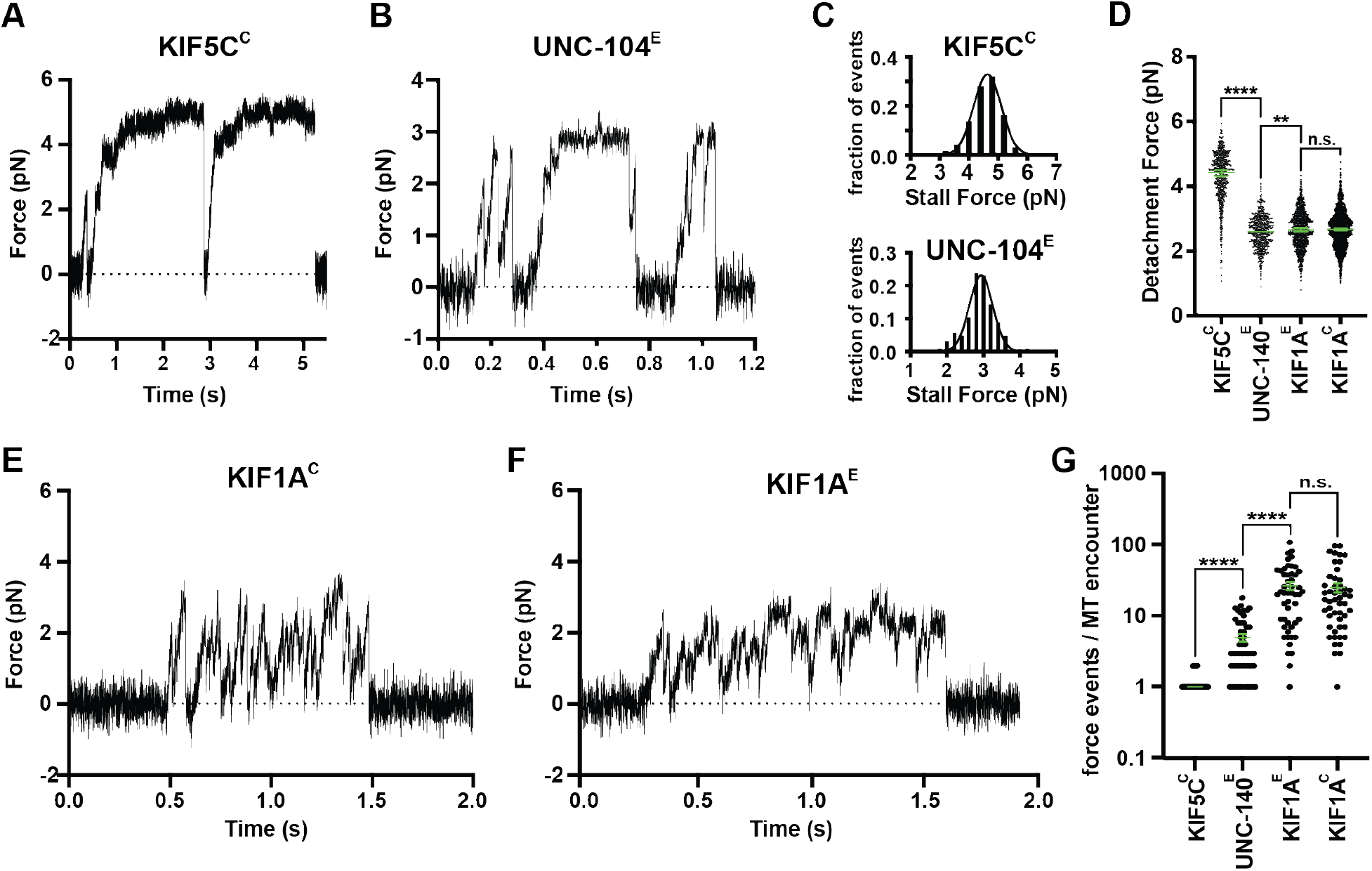
KIF1A and UNC-104 detach under low force and rapidly reattach to the MT. (A,B,E,F) Representative force vs. time records of bead movement driven by single molecules of (A) kinesin-1 KIF5C(1-560) in COS-7 cell lysates (KIF5CC), (B) kinesin-3 UNC-104(1-389) purified from *E. coli* bacteria (UNC-104^E^), (E) kinesin-3 KIF1A(1-393)-LZ in COS-7 cell lysates (KIF1A^C^), and (F) kinesin-3 KIF1A(1-393)-LZ purified from *E. coli* bacteria (KIF1A^E^). (C) Stall force histograms of KIF5CC (4.64 ± 0.01 pN, mean ± SEM from Gaussian fit; stall plateaus ≥200 ms; *N* = 197) and UNC-104^E^ (2.94 ± 0.03 pN, stall plateaus ≥10 ms; *N* = 126) compiling forces at *k* = 0.05 – 0.06 pN/nm. (D) Detachment forces. Green bars indicate the median values with quartiles. KIF5CC: 4.43 [3.79, 4.86] pN, *N* = 557; UNC-104^E^: 2.59 [2.23, 2.94] pN, *N* = 561; KIF1A^E^: 2.65 [2.25, 3.05] pN, *N* = 1044; KIF1A^C^: 2.66 [2.25, 3.01] pN, *N* = 1912. (G) Number of motor engagement events per MT encounter. KIF5CC: 1.2 ± 0.1 (mean ± SEM), *N* = 50; UNC-104^E^: 5.0 ± 0.6, *N* = 50; KIF1A^E^: 24 ± 3, *N* = 50; KIF1A^C^: 25 ± 4, *N* = 50.

For UNC-104, we purified a truncated [UNC-104(1-389)] and biotinylated version from *E. coli* cells (UNC-104^E^) (42). Individual UNC-104^E^ motors were processive in the absence of load (Fig S2A), and frequently detached under load before reaching a stall plateau (Fig 1B). UNC-104^E^ motors stalled (≥10 ms criterion) at 3 ± 0.6 pN (±SEM) (Fig 1C) but frequently detached before stalling at an average detachment force of 2.6 pN (Fig 1D, Table 1). Interestingly, unlike kinesin-1 KIF5CC, UNC-104^E^ motors frequently reengaged with the MT track: UNC-104^E^ produced 5.0 ± 0.6 events (±SEM) per MT encounter whereas KIF5CC exhibited only 1.2 ± 0.1 events (Fig 1G).

For KIF1A, we used a truncated version that is constitutively active [KIF1A(1-393)] followed by a leucine zipper (LZ) to ensure the motor is in a dimeric state (16), and compared the behavior of KIF1A(1-393)-LZ motors present in COS-7 cell lysates (KIF1A^C^) to those expressed and purified from *E. coli* bacteria (KIF1A^E^). Individual KIF1A motors underwent fast motility in the absence of load (Figs S2 and 6) but in contrast to KIF5C and UNC-104, KIF1A motors did not exhibit motor stalling; rather, KIF1A motors rapidly detached from the MT when subjected to force (Fig 1E&F). We measured an average detachment force of 2.7 pN for KIF1A motors expressed in mammalian or bacterial cells (Fig 1D, Table 1). Interestingly, KIF1A motors quickly rebound to the MT after detachment and moved forward again, presumably due to the motor’s high on-rate towards MTs (48). These rapid detachment and reattachment cycles result in a “clustering” of force generation events (Fig 1E&F). The average number of rebinding events per MT encounter is even higher for KIF1A than UNC-104 at 24 ± 3 (±SEM) events for KIF1A^E^ and 25 ± 4 events for KIF1A^C^ (Fig 1G).

**Table 1.** Single-molecule detachment forces.

| Kinesin motor | COS-7 cell lysate | *E. coli* expressed |
| --- | --- | --- |
| kinesin-1 KIF5C | 4.4 [3.8, 4.9] pN | ND |
| kinesin-3 UNC-104 | ND | 2.6 [2.2, 2.9] pN |
| kinesin-3 KIF1A | 2.7 [2.3, 3.0] pN | 2.7 [2.3, 3.1] pN |
| kinesin-3 KIF1A-V8M | 2.0 [1.7, 2.2] pN | 1.9 [1.7, 2.2] pN |
| kinesin-3 KIF1A-Y89D | 1.0 [0.9, 1.2] pN | 1.0 [0.9, 1.2] pN |

Data reported as Mean [quartiles]. ND, not determined.

### KIF1A disease variants are predicted to impact motor force generation

We hypothesized that the mechanism of KIF1A force generation is similar to that of kinesin-1 and uses nucleotide-dependent conformational changes of the NL to facilitate force generation. To test this, we looked for KAND-associated mutations located in regions predicated to be critical for NL docking. We mapped KAND-associated mutations onto the protein sequence (Fig 2A, red lines) and structure (Fig 2B, red circles) of the KIF1A motor domain [PDB 4UY0, (30)]. The majority (21/31) of KAND-associated mutations cluster within functional elements critical for MT binding, nucleotide binding, or force generation (Fig 2A&B, Table 2). In particular, two *de novo* KAND-associated mutations, V8M and Y89D, are located in elements important for kinesin-1 motors to step against force (Fig 2C). To delineate the local and global effects of these mutations on the KIF1A motor domain, we performed all-atom MD simulations of WT or mutant motor domains interacting with the MT in their ATP-bound state [post-power stroke, PDB 4UXP, (30)]. Four replicate simulations of at least 200 ns each were carried out and analysis across replicate simulations was used to predict statistically significant differences in residue-residue distances between WT KIF1A and the KAND mutant motors (p<10^-5^, V8M Fig 3; Y89D Fig 4).

**Fig 2.**
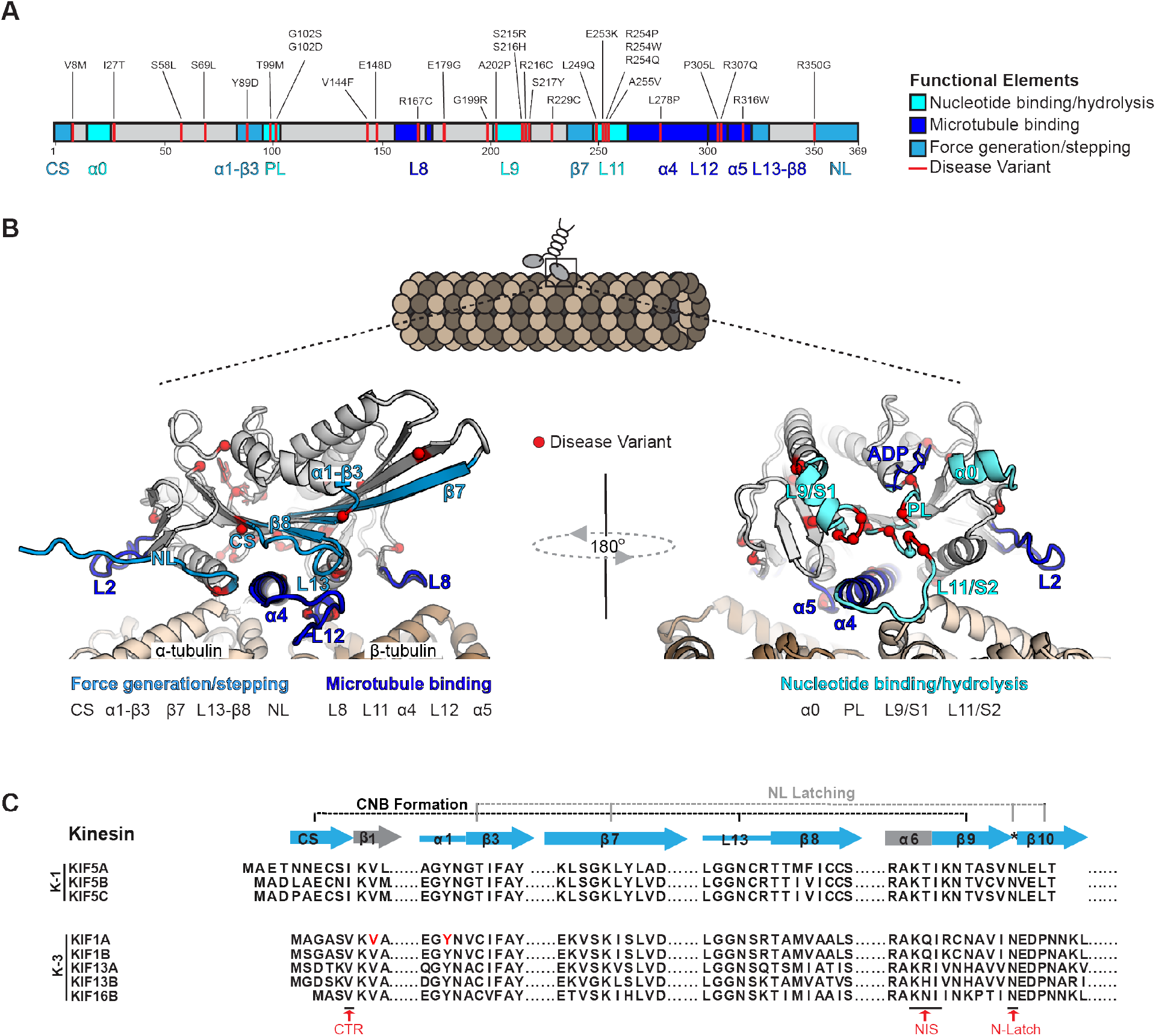
KIF1A disease variants cluster within regions of the motor domain critical for MT binding, nucleotide binding/hydrolysis, and stepping/force generation. (A,B) KIF1A disease variants (red) mapped onto the KIF1A motor domain (A) protein sequence and (B) ribbon representation of the ADP-bound, tubulin-bound state (PDB 4UYO). Functional elements are indicated as dark blue: MT binding (Loop8, α4, Loop12, α5); medium blue: stepping/force generation (CS, α1-β3, β8, Loop 13, NL); and cyan: nucleotide binding/hydrolysis (Loop 9, Loop 11, P Loop, α0). (C) Alignment of sequences implicated in force generation for the human kinesin motor domain from the kinesin-1 and kinesin-3 families.

**Fig 3.**
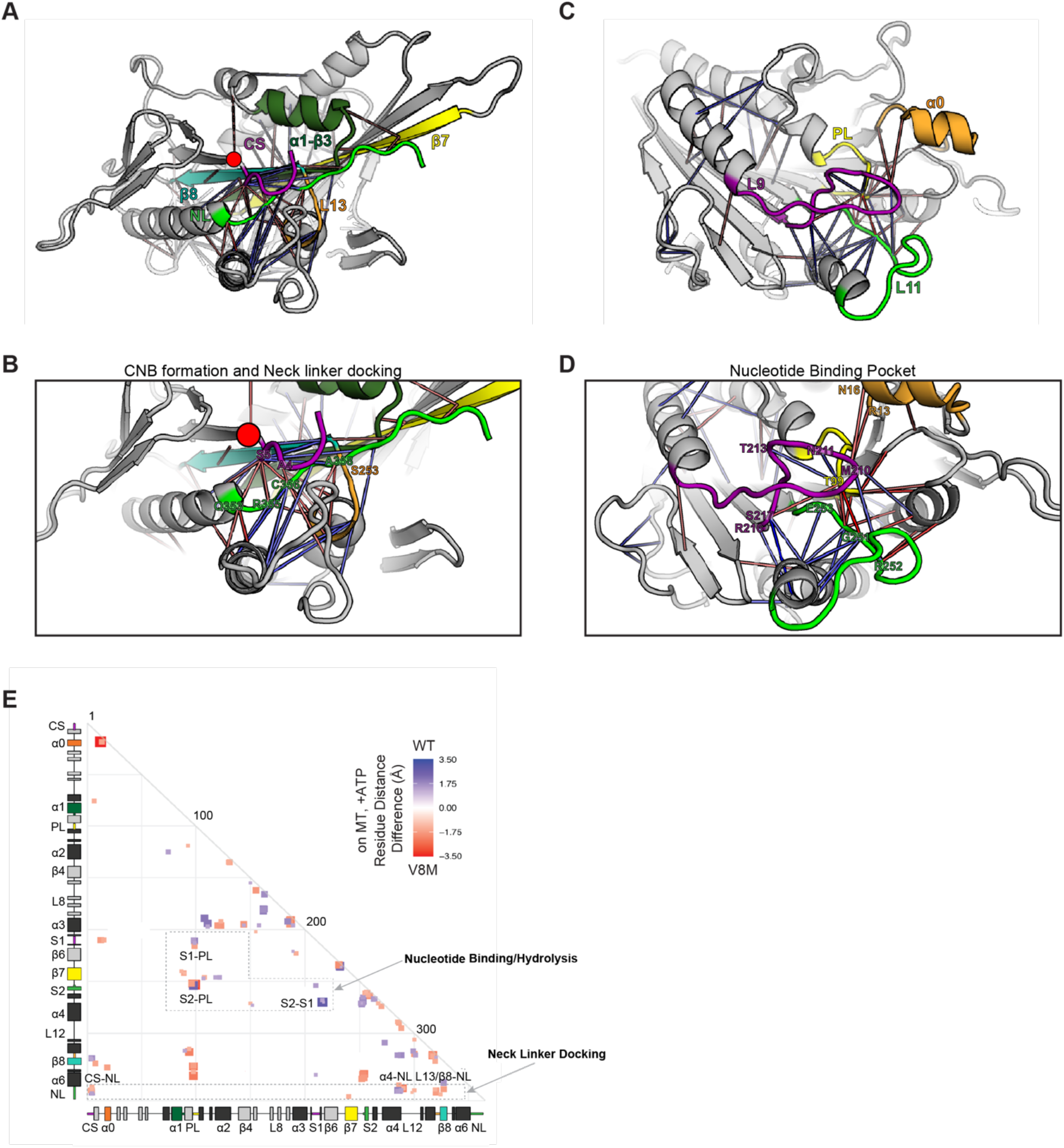
MD simulations predict that the V8M mutation alters NL docking and catalytic site closure. (A-D) Ribbon representation of the KIF1A motor domain in the ATP-bound, tubulin-bound state (PDB 4UXP). The V8M mutation (β1) is denoted as a red circle. Red lines depict residue-residue distances that are shorter in the V8M mutant whereas blue lines depict residue-residue distances that are shorter in the WT motor. The magnitude of the distance change is indicated by color intensity. (A,B) View of the NL docking pocket. In this post-power stroke state, the NL (green) is docked along the core motor domain. Secondary structures are indicated as purple: CS; dark green: α1-β3; yellow: β7; teal: β8; and orange: Loop 13 (L13). (C,D). View of the nucleotide-binding pocket. Secondary structures are indicated as purple: Loop 9/Switch1 (L9/S1); green: Loop 11/Switch2 (L11/S2); yellow: P Loop (PL); and orange: α0. (E) Differences in residue-residue distances between WT KIF1A and V8M mutant motor in the ATP-bound, tubulin-bound state determined in MD simulations. The secondary structure elements are laid out along the x- and y-axes with α-helices in black and β-strands in grey or colored according to (A). Residue-residue interactions that are significantly (p<10^-5^) shorter in V8M mutant (red) or the WT motor (blue) are displayed on the grid. The magnitude of the distance change is indicated by color intensity.

**Fig 4.**
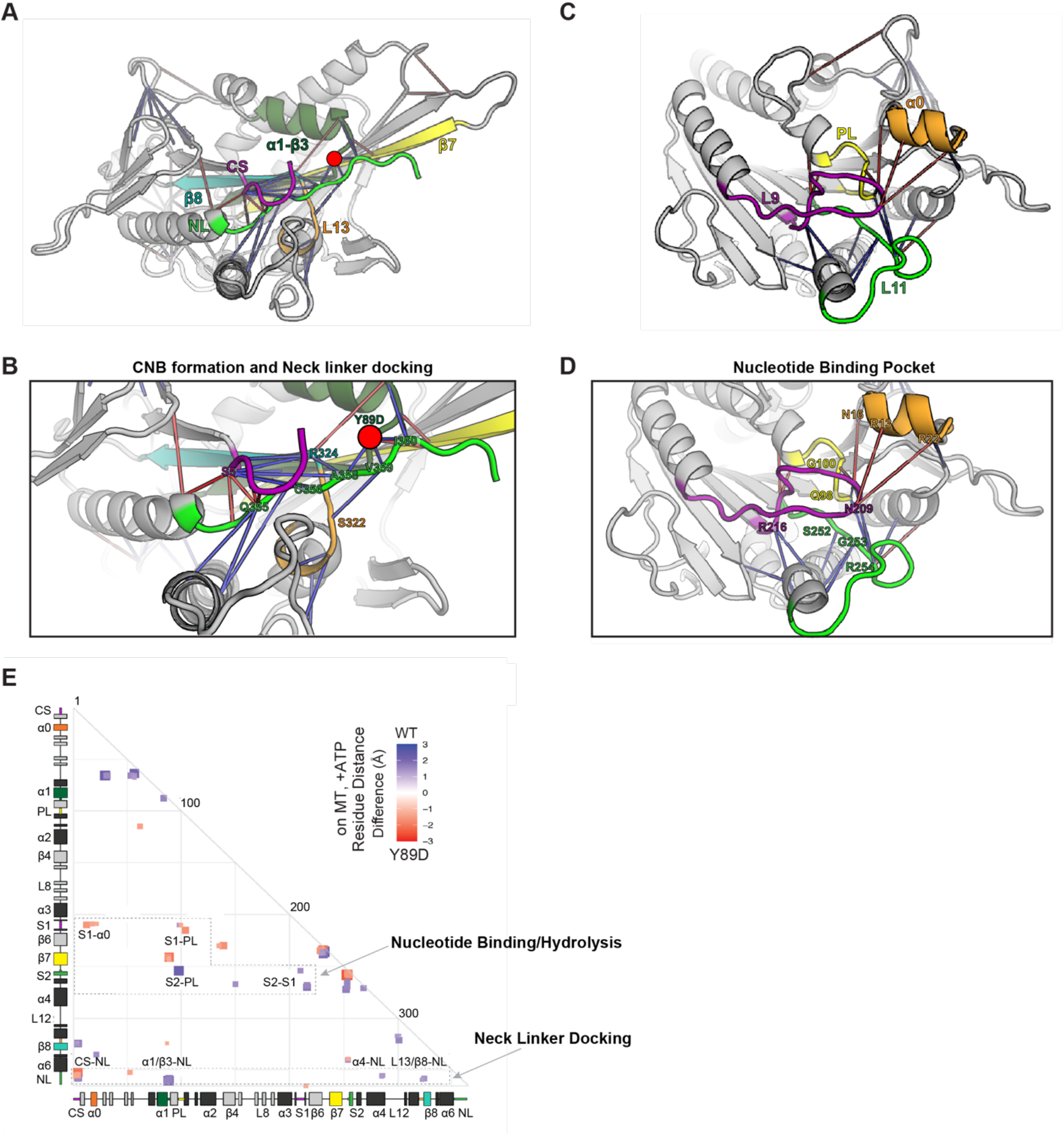
MD simulations predict that the Y89D mutation alters NL docking and catalytic site closure. (A-D) Ribbon representation of the KIF1A motor domain in the ATP-bound, tubulin-bound state (PDB 4UXP). The Y89D mutation (α1-β3) is denoted as a red circle. Red lines depict residue-residue distances that are shorter in the Y89D mutant whereas blue lines depict residue-residue distances that are shorter in the WT motor. The magnitude of the distance change is indicated by color intensity. (A,B) View of the NL docking pocket. In this post-power stroke state, the NL (green) is docked along the core motor domain. Secondary structures are indicated as purple: CS; dark green: α1-β3; yellow: β7; teal: β8; and orange: Loop 13 (L13). (C,D) View of the nucleotide-binding pocket. Secondary structures are indicated as purple: Loop 9/Switch1 (L9/S1); green: Loop 11/Switch2 (L11/S2); yellow: P Loop (PL); and orange: α0. (E) Differences in residue-residue distances between WT KIF1A and the Y89D mutant motor in the ATP-bound, tubulin-bound state determined in MD simulations. The secondary structure elements are laid out along the x-and y-axes with a-helices in black and β-strands in grey or colored according to (A). Residue-residue interactions that are significantly (p<10^-5^) shorter in Y89D mutant (red) or the WT motor (blue) are displayed on the grid. The magnitude of the distance change is indicated by color intensity.

For the V8M mutation, the MD simulations predict local changes in residue-residue interactions important for NL-dependent motor stepping and force generation (Fig 3A,B,E). Enhanced interactions are observed between the initial residues of β9 of the NL and the second residue (S6) of the CS (Fig 3A&B, red lines; Fig 3E, red box marked CS-NL), which may contribute to CNB formation and force output. However, reduced interactions are observed for the remainder of β9 and elements that position it for NL docking. In particular, reduced interactions are observed between β9 and residues of α4 that make up the docking pocket (Fig 3A&B, blue lines; Fig 3E, blue box marked α4-NL). Thus, the V8M mutation may position the CS such that it sterically occludes the NL’s access to the docking pocket. The MD simulations also predict reduced interactions between elements important for coordinating and hydrolyzing nucleotide (Fig 3C&D, blue lines; Fig 3E, boxes marked S1-PL and S2-S1). As closure of the switch regions is necessary for ATP hydrolysis (26, 49, 50), these results suggest that the V8M mutant motor may have problems coordinating and/or hydrolyzing ATP and therefore have a reduced velocity compared to WT motors.

For the Y89D mutation, the MD simulations predict more severe restrictions on NL docking and thus a greater impact on motor stepping and force generation. Specifically, the MD simulations reveal reduced interactions important for positioning β9 of the NL in the α4-lined docking pocket (Fig 4A&B, blue lines; Fig 4E, blue box marked α4-NL) and for subsequent docking of β10 along the core motor domain (Fig 4&B, blue lines; Fig 4E, blue boxes marked α1/β3-NL and L13/β8-NL). In addition, the MD simulations revealed mixed effects of the Y89D mutation on interactions between elements in the nucleotide-binding pocket. There are enhanced interactions between elements important for gating and capture of nucleotide (Fig 4C&D, red lines; Fig 4E, red boxes marked S1-α0) as well as reduced interactions between elements important for nucleotide hydrolysis and exchange (Fig 4C&D, blue lines; Fig 4E, blue boxes marked S2-PL and S2-S1) (26, 49-51). Therefore, these results suggest that although the mutant motor may have no restrictions on binding ATP, it may display a reduced ability to hydrolyze ATP and undergo processive motility.

**Table 2.** KAND-associated mutations that map to the KIF1A motor domain.

| KIF1A functional region | KAND-associated mutations |
| --- | --- |
| MT binding | $\alpha 4$: L278P<br>Loop12: P305L and R307Q<br>$\alpha 5$: R316W<br>$\alpha 6$: R350G |
| nucleotide binding and hydrolysis | P Loop: T99M and 102S/D<br>Loop 9 (Switch 1): A202P, S215R/H, R216C, S217Y<br>Loop 11 (Switch 2): L249Q, E253K, R254P/W/Q, A255V |
| force generation and motor stepping | $\beta 1$: V8M<br>$\alpha 1$-$\beta 3$: Y89D |

### Impact of V8M and Y89D mutations on the force generation of homodimeric motors

To examine the effects of the V8M and Y89D mutations on the force output of the motors, we used optical tweezers and motors attached to beads under single-molecule conditions as done for the WT motor. Biotinylated KIF1A-AviTag motors containing the V8M or Y89D mutations were bound to streptavidin-coated trapping beads from lysates of COS-7 cells (KIF1A^C^-V8M/Y89D) or after purification from *E. coli* bacteria (KIF1A^E^-V8M/Y89D). Both mutant motors were sensitive to small opposing forces exerted by the trap. Similar to the WT motor, the V8M and Y89D motors detached from the MT before reaching a stall plateau (Fig 5A-D). However, both mutant motors displayed an impaired force output as their average detachment forces (1.9 and 1.0 pN, respectively) were significantly reduced (Fig 5E&F; Table 1) compared to WT KIF1A. The reduced force output of the mutant motors is consistent with our MD simulations that predict that the KAND mutations would impair docking of β9 and/or β10 of the NL to the core motor domain (Figs 3 and 4). Interestingly, similar to the WT motor, the mutant motors quickly rebound to the MT after detaching (Fig 5A-D), resulting in a clustering of force-generating events.

**Fig 5.**
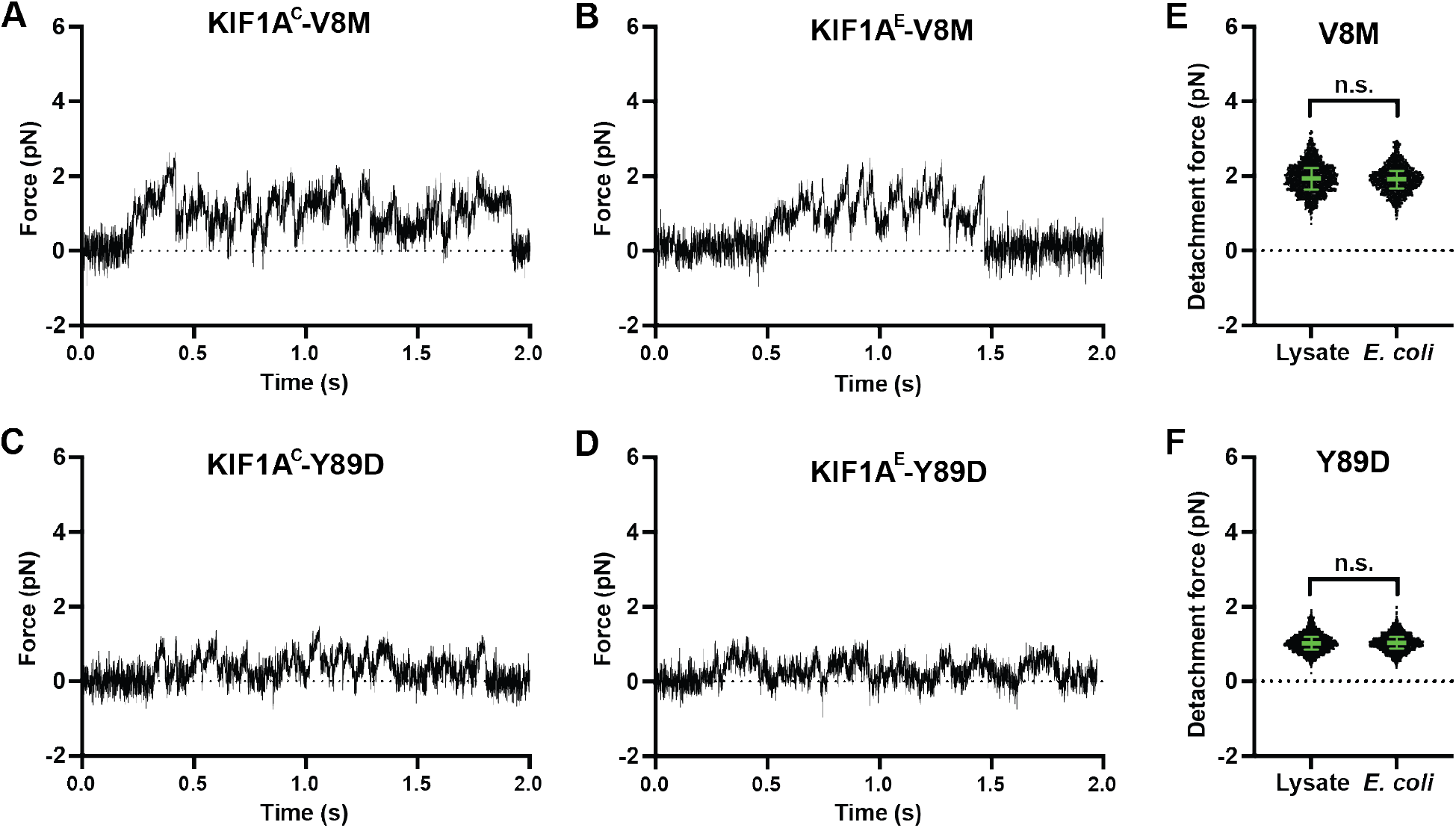
V8M and Y89D mutations result in decreased force output for KIF1A motors. (A-D) Representative force vs. time records of bead movement driven by single molecules of KIF1A-V8M mutant motors in (A) COS-7 cell lysates (KIF1A^C^-V8M) or (B) purified from *E. coli* (KIF1A^E^-V8M) or KIF1A-Y89D mutant motors in (C) COS-7 cell lysates or (D) purified from *E. coli.* (E,F) Detachment forces of (E) V8M and (F) Y89D mutant motors. Green bars indicate the median values with quartiles. V8M: 1.94 [1.65, 2.22] pN, *N* = 1343; Y89D: 1.02 [0.87, 1.19] pN, *N* = 1468.

### Impact of V8M and Y89D mutations on unloaded motility properties of homodimeric motors

We next used fluorescence-based single-molecule motility assays to examine the behavior of WT or KAND mutant KIF1A motors under unloaded conditions. For this work, we used KIF1A(1-393)-LZ motors tagged with a HaloTag^®^ (for fluorescent labeling with JF552 ligand) and a FLAG tag and present in COS-7 cell lysates (Fig 6). The motors were added to flow chambers containing polymerized MTs and their single-molecule motility properties were examined using total internal reflection florescence (TIRF) microscopy (52). As expected, the truncated, homodimeric WT motor displayed fast [2.1 ± 0.1 μm/s (mean ± SEM), Fig 6C] and super-processive (16.7 [10.2, 27.2] μm (median [25%, 75%]), Fig 6D) motility with a high landing rate of 0.22 ± 0.01 events·nm^-1^onM^-1^os^-1^ (mean ± SEM; Fig 6E, Table 3), consistent with previous work (18, 48). The homodimeric V8M mutant motors displayed a significant decrease in overall velocity (1.3 ± 0.1 μm/s, Fig 6C), processivity (4.1 [2.1, 7.0] μm, Fig 6D), and landing rate (0.05 ± 0.01 eventsonm^-1^onM^-1^os^-1^, Fig 6E) (Table 3). The reduced velocity of the V8M mutant motors is consistent with the MD simulations that predict allosteric effects on the nucleotide-binding pocket that result in reduced catalytic site closure and reduced ATP hydrolysis (Fig 3C-E).

**Fig 6.**
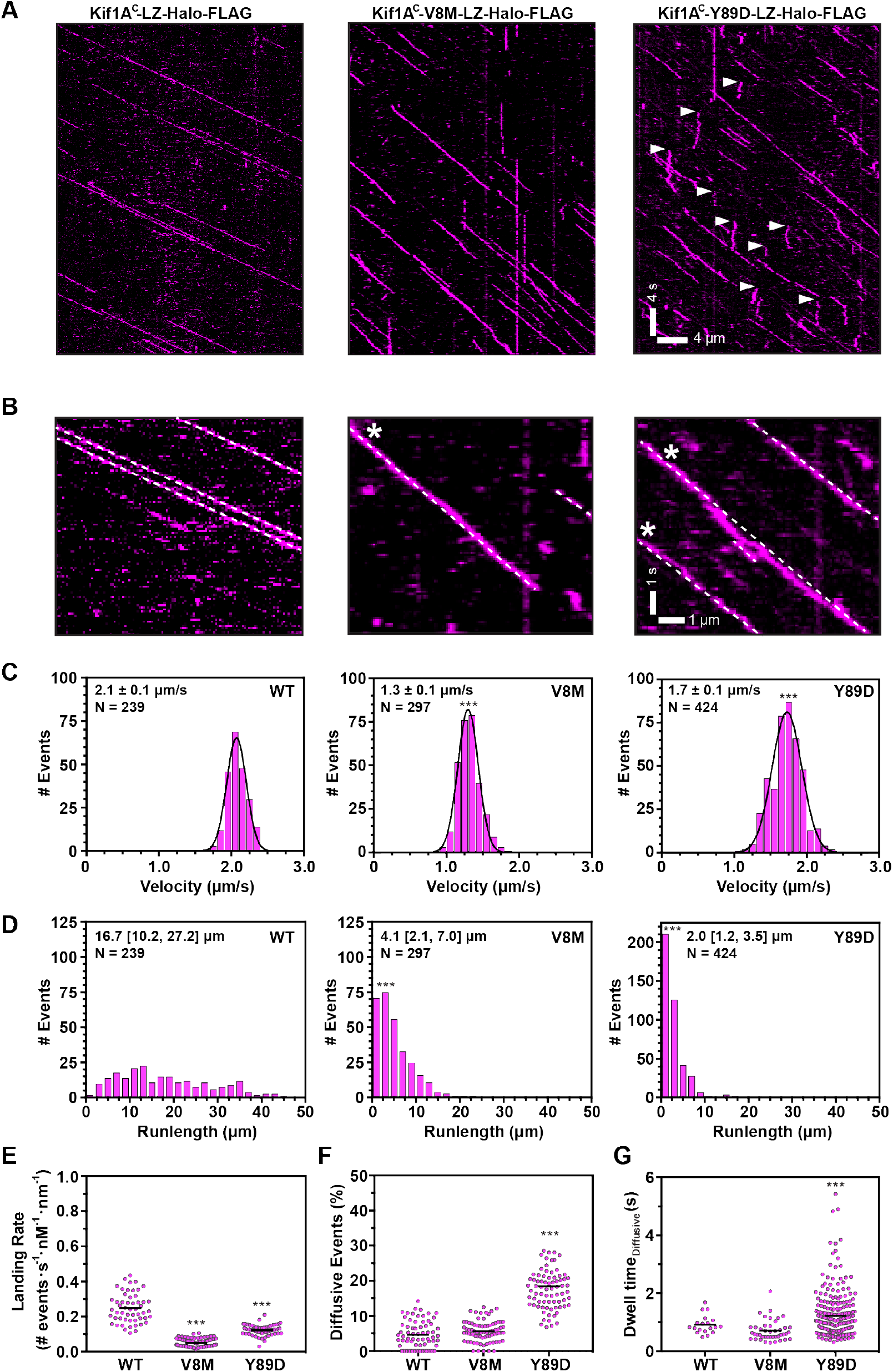
Homodimeric V8M and Y89D mutant motors display decreased speed, processivity, and landing rates. WT or mutant motors fused to a HaloTag^®^ and a FLAG-tag, and fluorescently labeled with the JF552-HaloTag ligand, were analyzed in standard single-molecule motility assays using TIRF microscopy. (A) Representative kymographs with time displayed on the y-axis (bar, 4 s) and distance displayed on the x-axis (bar, 4 μm). White arrowheads indicate motility events scored as diffusive events. (B) Magnified view of the kymographs in (A) with x-axis bar, 1 μm and y-axis bar, 1 s. Dotted white lines indicate linear motility; white asterisks indicate “wobbly” events that deviate from linear motility. (C-E) Quantification of motility properties. From the kymographs, single-motor (C) velocities (mean ± SEM), (D) run lengths (median [quartiles]), and (E) landing rates were determined. For (E), each dot indicates the motor landing rate along a single MT with the population mean indicated by a horizontal black line across three independent experiments for each construct; ***, p<0.001 as compared to the WT motor. (F,G) Quantification of diffusive motility events. From the kymographs, the (F) percentage of diffusive events (dwell time longer than 400 ms, net displacement less than 200 μm) and (G) dwell time of the diffusive events were determined. Each spot indicates the behavior of a single motor. Black horizontal lines indicate the population mean; ***, p<0.001 as compared to WT motor.

The Y89D mutant motors also displayed a decrease in velocity (1.7 ± 0.1 μm/s, Fig 6C), processivity (2.0 [1.2, 3.5] μm, Fig 6D), and landing rate (0.12 ± 0.01 eventsonm^-1^onM^-1^os^-1^, Fig 6E) as compared to the WT motor (Fig 6C-E, Table 3). Further examination of the kymographs indicated two additional differences in the motility behavior of Y89D mutant motors. First, the tracks of Y89D motility were not smooth but rather the motors appeared to “wobble” or move sideways as they walked along the MT track (Fig 6B). Second, a large number of non-productive, diffusive events (net displacement along the MT < 200 nm) were observed (Fig 6A, white arrowheads). We quantified the percentage of diffusive events with a dwell time greater than 400 ms for each motor (Fig 6F). The Y89D mutant motors displayed a greater percentage of diffusive events (18.5 ± 1.2% of binding events) than the WT or V8M motors (4.4 ± 0.5% and 5.7 ± 0.3 %, respectively) (Fig 6F, Table 3) and the duration of the diffusive events was longer for the Y89D mutant motors (1.3 ± 0.2 s) than for the WT or V8M mutant motors (0.81 ± 0.01 s and 0.69 ± 0.02 s, respectively, Fig 6G). The increase in diffusive events suggests that the Y89D mutant motor often engages in a weak MT-binding state.

To ensure that the changes in motility of the V8M and Y89D motors were due to direct effects on motor behavior rather than indirect alterations in the cell lysate context, we purified SNAP- and His-tagged homodimeric WT, V8M, and Y89D motors from *E. coli* bacteria. The purified recombinant WT motors displayed fast (2.5 ± 0.2 μm/s, Fig S2B&C) and superprocessive (12.2 [6.7,18.4] μm, Fig S2D) motility (Table 3). Similar to the effects observed for the mammalian-expressed mutant motors, the recombinant KIF1A-V8M^E^ and KIF1A-Y89D^E^ mutant motors were slower (1.3 ± 0.1 μm/s and 1.7 ± 0.2 μm/s, respectively, Fig S2F&I) and displayed a reduced processivity (7.3 [4.4,12.2] μm and 6.3 [4.0, 7.0] μm, respectively, Fig S2F,I) as compared to the WT^E^ motor (Table 3). Overall, we conclude that as homodimeric motors, the V8M motor shows a significant impairment in velocity and landing rate whereas the Y89D motor shows a significant impairment in processivity and in its ability to engage in processive rather than diffusive motility (Table 3).

**Table 3.** Unloaded Single-Molecule Motility Properties.

| KIF1A motor | velocity<br>($\mu\text{m}/\text{sec}$)* | run length<br>($\mu\text{m}$)<sup>#</sup> | landing rate<br>(# events $\cdot \text{s}^{-1} \cdot \text{nM}^{-1} \cdot \text{nm}^{-1}$) | % diffusive events<br>(diffusive / total events) |
| --- | --- | --- | --- | --- |
| WT/WT<sup>C</sup> | $2.1 \pm 0.1$ | 16.7 [10.2,27.2] | $0.22 \pm 0.01$ | $4.4 \pm 0.5$ |
| V8M/V8M<sup>C</sup> | $1.3 \pm 0.1$ | 4.1 [2.1,7.0] | $0.05 \pm 0.01$ | $5.7 \pm 0.3$ |
| Y89D/Y89D<sup>C</sup> | $1.7 \pm 0.1$ | 2.0 [1.2, 3.5] | $0.12 \pm 0.01$ | $18.5 \pm 1.2$ |
| WT/V8M<sup>C</sup> | $1.3 \pm 0.1$ | 10.0 [5.5, 15.0] | ND | ND |
| WT/Y89D<sup>C</sup> | $1.9 \pm 0.1$ | 9.0 [4.9, 16.6] | ND | ND |
| WT/WT<sup>E</sup> | $2.5 \pm 0.2$ | 12.2 [6.7,18.4] | ND | ND |
| V8M/V8M<sup>E</sup> | $1.3 \pm 0.1$ | 7.3 [4.4,12.2] | ND | ND |
| Y89D/Y89D<sup>E</sup> | $1.7 \pm 0.2$ | 6.3 [4.0, 7.0] | ND | ND |

\*Data reported as Mean ± SEM. <sup>#</sup> Data reported as Median [quartiles]. ND, not determined.

### Impact of V8M and Y89D mutations on unloaded motility properties of heterodimeric motors

The V8M and Y89D mutations found in KAND patients are inherited in an autosomal dominant manner, indicating that the disease allele can influence transport even in the presence of a WT allele. We thus examined the effect of the KAND mutations in the heterodimeric state where one motor domain is WT and the second motor domain harbors a KAND mutation. We tested several strategies for generating heterodimeric motors but were unable to achieve complete heterodimer formation (Fig S3). We thus co-transfected COS-7 cells with plasmids for expression of WT motors tagged with mNeonGreen (mNG) and KAND mutant motors tagged with HaloTag and FLAG tag (Fig 7A). We tested several imaging conditions to avoid artifacts related to either of the tags (Fig S4) and carried out single-molecule motility assays using TIRF microscopy. From the kymographs, motility events of heterodimeric motors were scored as co-motility in both the mNG and Halo(JF552) fluorescence channels (Fig 7B, E&H, left panels). To better visualize heterodimeric motor events, cartoon kymographs were generated to display motile events (Fig 7B, E&H, middle panels) and diffusive events (Fig 7B, E&H, right panels).

**Fig 7.**
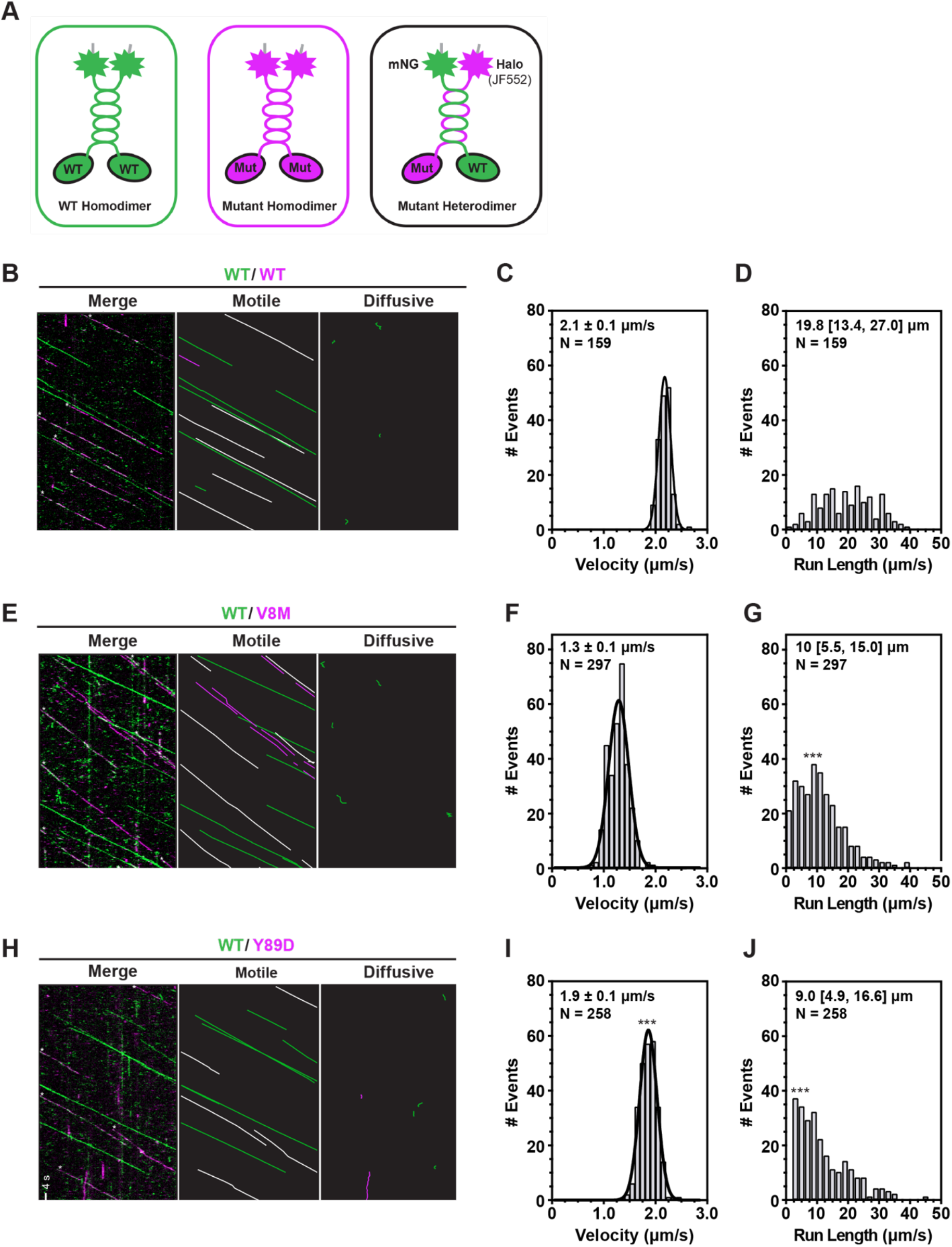
Heterodimeric WT/V8M and WT/Y89D mutant motors display decreased processivity. (A) Schematic of strategy for generating heterodimeric WT/mutant KIF1A motors. WT KIF1A(1-393)-LZ was tagged with monomeric NeonGreen (mNG) whereas V8M and Y89D motors were tagged with Halo-FLAG and labelled with JF552. When co-expressed in cells, three populations of motors are expected in TIRF single-molecule assays: homodimeric WT motors tagged with mNG, homodimeric mutant motors tagged with Halo/JF552, and heterodimeric WT/mutant motors tagged with mNG and Halo/JF552. (B-J) Single-molecule motility analyses based on TIRF microscopy data. (B,E,H) Representative kymographs (left) with time displayed on the y-axis (bar, 4 s) and distance displayed on the x-axis (bar, 4 μm). Cartoon kymographs were generated from merged kymograph to more clearly illustrate motile (middle) and diffusive events (right). From the kymographs, single-motor (C,F,I) velocities and (D,G,J) run lengths were determined and the data for each population were plotted as a histogram. The mean values ± SEM (for velocities) and median with quartiles (for run lengths) are indicated on each graph; ***, p<0.001 as compared to the WT motor.

The velocity (2.1 ± 0.1 μm/s) and run length (19.8 [13.4, 27.0] μm) of WT/WTC motors tagged with both mNG and Halo(JF552) fluorophores (Fig 7B,C&D) are comparable to those of KIF1A(393)-LZ-Halo-Flag motors (Fig 6; Table 3). The presence of the V8M motor domain resulted in a significant reduction in velocity (1.3 ± 0.1 μm/s Fig 7E,F) such that the heterodimeric WT/V8MC motor’s velocity is comparable to that of homodimeric V8M/V8MC mutant motors (Table 3). In addition, the processivity of WT/V8MC motors (10.0 [5.5, 15.0] μm, Fig 7E,G) was significantly reduced compared to WT/WT^C^ motors but was not as severely hindered as in the V8M/V8MC motors (Table 3).

The presence of the Y89D motor domain had minimal effects on velocity in the context of the heterodimeric WT/Y89DC motor (1.9 ± 0.1 μm/s, Fig 7H,I) as compared to the WT/WT^C^ motor but resulted in a significant reduction in the processivity (9.0 [4.9, 16.6] μm, Fig 7H,J) although these effects were not as severe as observed for the Y89D/Y89D homodimeric motors (Table 3). In addition, the WT/Y89DC heterodimeric motors did not exhibit the diffusive behavior of the Y89D/Y89DC homodimeric motors (Fig 7H). Collectively, these results suggest that when paired with a WT motor domain in a heterodimeric motor, both the V8M and Y89D mutations cripple the overall motility with greater effects on motor processivity than motor speed.

### Impact of V8M and Y89D mutations on transport of membrane-bound organelles in cells

We next sought to test whether these mutations impacted the ability of motors to work as a team to drive cargo transport in cells. To do this, we used an inducible recruitment strategy (53, 54) to link teams of motors to the surface of membrane-bound organelles and monitored their ability to drive organelle transport to the cell periphery (Fig 8A). To assess how teams of WT or KAND mutant KIF1A motors drive the transport of a low-load, membrane-bound organelle (36, 54, 55), motors were recruited to the surface of peroxisomes, and transport of peroxisomes to the cell periphery was assessed after 5, 10, or 30 minutes. Cargo location before and after motor recruitment was qualitatively scored as clustered (black), partially dispersed (dark grey), diffusively dispersed (light grey), or peripherally dispersed (white; Fig 8C).

**Fig 8.**
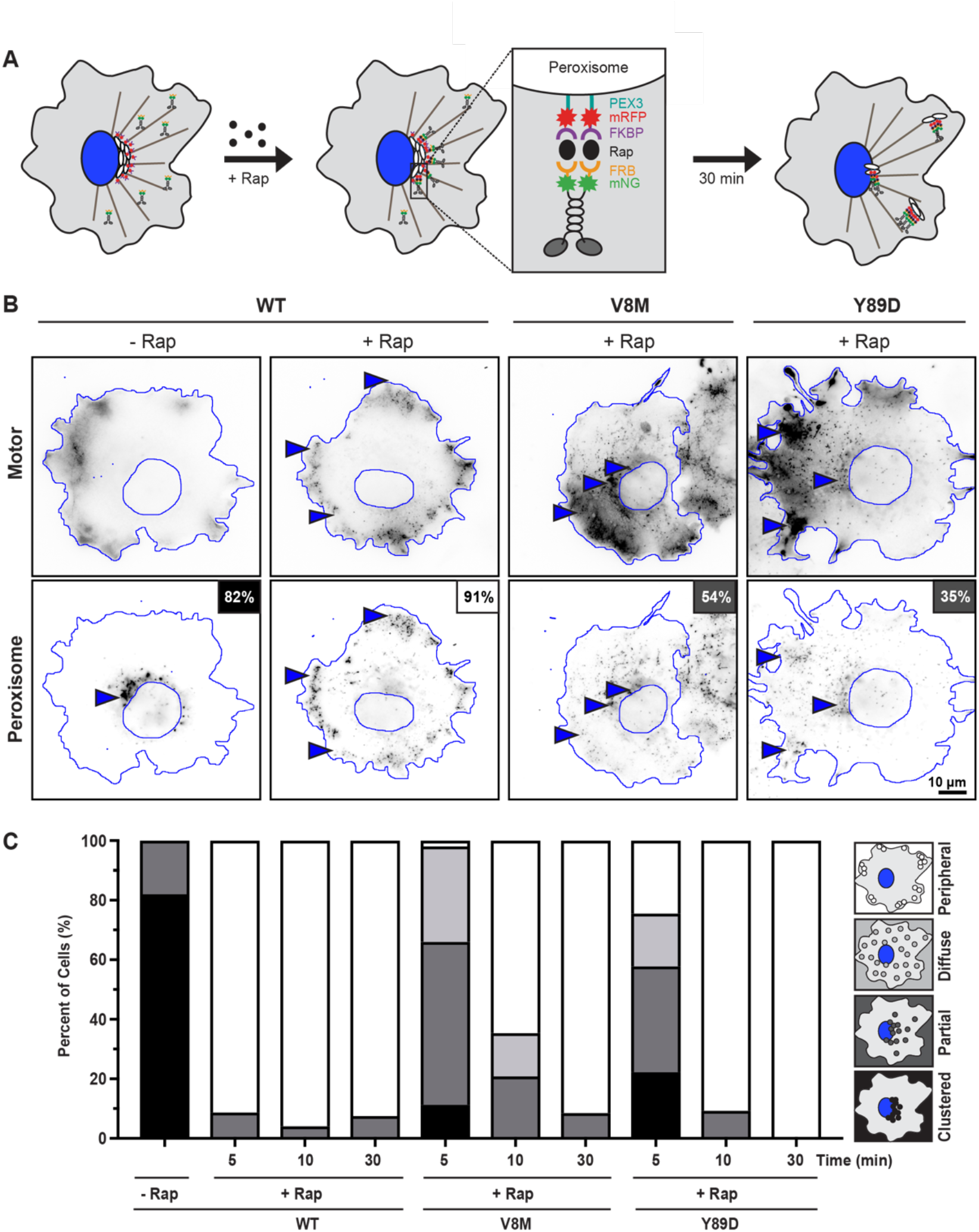
V8M and Y89D mutant motors show delayed transport of membrane-bound cargo in cells. (A) Schematic of the inducible motor recruitment assay. A kinesin motor fused to mNG and an FRB domain (KIF1A-mNG-FRB) is coexpressed in COS-7 cells with a peroxisome-targeting sequence (PEX3) fused to monomeric red fluorescent protein (mRFP) and an FKBP domain (PEX-mRFP-FKBP). Addition of rapamycin (+Rap) causes heterodimerization of the FRB and FKBP domains and recruitment of motors to the peroxisome membrane. Recruitment of active motors drives peroxisome dispersion to the cell periphery. (B) Representative images of peroxisome dispersion before (-Rap) and after (+Rap) recruitment of WT or mutant motors to the peroxisome surface. Blue lines indicate the nucleus and periphery of each cell. Blue arrowheads indicate peroxisomes. Scale bar, 10 μm. Percentages in the upper right corner indicate the percent of cells with the indicated dispersion phenotype: black: clustered peroxisomes; dark gray: partially dispersed peroxisomes; light gray: diffusely dispersed peroxisomes; white: peripherally dispersed peroxisomes. (C) Qualitative analysis of peroxisome dispersion. Cells were scored as clustered (black), partially dispersed (dark grey), diffusely dispersed (light grey), or peripherally dispersed (white). The phenotypes of *N* ≥ 43 cells across three experiments were combined into a stacked bar plot for each construct at each time point.

COS7 cells were co-transfected with a plasmid for the expression of WT or KAND mutant KIF1A(393)-LZ motors tagged with mNG and FRB domains and a plasmid for the expression of a peroxisome-targeted PEX-mRFP-FKBP fusion protein. In the absence of rapamycin, the PEX-RFP-FKBP peroxisomes were largely clustered in the center of the cell (93% of cells had clustered peroxisomes; Fig 8B,C) whereas KIF1A(393)-LZ-mNG-FRB motors accumulated at the periphery of the cell (Fig 8B). Addition of rapamycin resulted in recruitment of motors to the peroxisome surface via dimerization of the FRB and FKBP domains and motor activity drove dispersion of peroxisomes to the cell periphery. 5 minutes after recruitment of WT motors, 91% of cells (42/46) had peroxisomes dispersed to the periphery of the cell (Fig 8B,C). In contrast, 5 minutes after recruitment of teams of V8M or Y89D mutant motors, the peroxisomes failed to reach the periphery of the cell. Rather, 54% (29/53) of V8M-expressing and 35% (16/45) of Y89D-expressing cells displayed only partial peroxisome dispersion (Fig 8B,C).

We hypothesized that the impaired motility and force-generation properties of the V8M and Y89D motors could be overcome if the motors were given more time to complete the transport event. We thus repeated the peroxisome dispersion assay but waited 10 or 30 minutes after recruitment of teams of V8M or Y89D mutant motors to assess peroxisome localization. At 10 min after rapamycin-induced motor recruitment, 65% (31/48) of cells expressing the V8M mutant motor and 91% (39/43) of cells expressing the Y89D mutant motor displayed peripheral dispersion of the peroxisomes as compared to 96% (47/49) of cells expressing the WT motor (Fig 8C). After 30 min of motor recruitment, the V8M and Y89D mutant motors were able to achieve peroxisome dispersion [91% (43/47) and 100% (49/49) of cells, respectively] to the same extent as the WT motor [92% (49/53) of cells, Fig 8C]. These results suggest that despite reduced force output, processivity, and velocity under single-molecule conditions, the V8M and Y89D mutant motors can drive cargo transport if the cargo imposes minimal load and the motors are given a longer time frame to complete the transport event.

## DISCUSSION

Kinesin-3 motors drive a large number of intracellular trafficking events yet their ability to generate and sustain force has not been investigated. We find that, unlike conventional kinesin-1, mammalian KIF1A motors and *C. elegans* UNC-104 motors detach from the MT track under low forces. Furthermore, both motors rapidly reattach to the MT and continue forward motion, a property that may enable fast transport of presynaptic vesicles over long distances. While KIF1A motors do not stall under load, UNC-104 motors can stall before detaching. We find that the disease-associated V8M and Y89D mutations compromise the force output of single motors and result in decreased velocity, processivity, and landing rate via allosteric effects on regions of the core motor domain responsible for NL docking and the coordination and binding of nucleotide.

The mutant motors also show a delay in their ability to transport a low-load cargo in cells. These results highlight the benefits of combining single-molecule assays with structure-based simulations to investigate how subtle sequence changes can impact the mechanical output of a motor.

### KIF1A readily detaches from microtubules under load but rapidly reattaches for persistent motility

Previous studies of kinesin-3 motors focused on their striking motility properties under no-load conditions. In these assays, truncated and thus constitutively-active versions of dimeric kinesin-3 motors were found to move with high speeds, to be superprocessive, and to readily engage with the MT (high landing rate) (16-18, 48, 56-58). Here we provide the first analysis of mammalian kinesin-3 motors under load and note several interesting aspects of KIF1A force generation that are likely to impact its cellular functions.

First, single KIF1A motors do not stall when subjected to resisting forces but rather, they rapidly detach from the MT track; this is in stark contrast to the ability of single kinesin-1 motors to resist detachment under load (19, 23, 24, 59). A high load-dependent detachment rate is consistent with previous work showing that kinesin-3 motors give up easily when forced to compete with kinesin-1 motors in driving cargo transport (39-41). Interestingly, kinesin-2 (KIF3A/KIF3B) and kinesin-5 (Eg5) motors also have a tendency to detach at moderate forces in optical trap assays (60-66) and to give up easily when in competition with kinesin-1 (40).

Second, KIF1A motors can only sustain an average 2.7 pN of force before detachment from the MT track; this is in stark contrast to the ability of kinesin-1 motors to sustain 4-6 pN of force (19, 24, 27, 36). It seems unlikely that the detachment of KIF1A at low forces is due to the strength of the motor-MT interaction as KIF1A has a higher MT affinity than kinesin-1 both in the ADP-bound (weak MT affinity) and the ATP-bound (strong MT affinity and force-bearing) states (30, 48). It seems more likely that the detachment of KIF1A at low forces can be attributed to a mechanical/structural feature of this motor. An intriguing possibility is that the length of the N-terminal extension that proceeds the CS impacts the strength of the CNB and thus the force output of the motor. Kinesin-3 motors lack an N-terminal extension (Fig S1) and recent structural studies and MD simulations of KIF13B showed that this kinesin-3 motor forms a short CNB with weaker CS-NL interactions than kinesin-1 (38, 67). At first glance, previous work on KIF1A’s *C. elegans* homolog, UNC-104, would appear to contradict this model as UNC-104, which also lacks an N-terminal CS extension, frequently countered forces up to 6 pN (42). However, we have recently determined that the forces measured with this optical tweezers setup were likely affected by an unintended electronic low-pass filtering of the trapping data so that the reported maximal force of 6 pN force is retrospectively estimated to be closer to 4 pN (24). Indeed, when we repeated the experiments with a modern optical tweezers setup, we found that UNC-104 stalls at 3 pN (Fig 1C). Thus, like KIF1A, single UNC-104 motors sustain lower forces than kinesin-1 motors.

Third, after detachment, KIF1A motors rapidly reattach to the MT and again move forward against the trap. This behavior is consistent with the role of the kinesin-3-specific K-loop (loop 12) whose positively-charged residues are responsible for the high landing rate of KIF1A motors (48, 56). We note that the rapid detachment and reattachment of single KIF1A motors results in a characteristic sawtooth pattern for the force vs. time plot that has not been observed for other motors to date.

### KAND Mutations Provide Insight into a Conserved Mechanism of Kinesin Force Generation

Recent structural and biochemical assays with dimeric kinesin-1 motors have provided strong support for the model that nucleotide-dependent conformational changes in the NL facilitate force generation. NL docking is initiated by an ATP-dependent conformational change in α6 that drives a two-step NL docking: zipping together of the NL’s β9 with the CS (β0) to form the CNB and then latching of the NL’s β10 along the surface of the core motor domain (27, 29, 36). Structural studies have shown that similar ATP-induced changes occur to α6 and the NL in members of the kinesin-3 and kinesin-5 families (30-35, 37), supporting the hypothesis that NL docking is a force-generating mechanism utilized by all superfamily members. Our work provides the first test of this model for a member of the kinesin-3 family.

We focused on two *de novo KIF1A* disease variants, V8M and Y89D, as these residues are predicted to have roles in CNB formation and NL docking based on their a) location in structural elements of the motor domain associated with force generation in kinesin-1 motors, and b) occurrence in residues that are highly conserved across the kinesin superfamily (Figs 2C and S1). Our MD simulations found that the V8M and Y89D mutations impair docking of the N-terminal (β9) or C-terminal (β10) portions of the NL to the KIF1A motor domain, respectively (Figs 2 and 3). Indeed, using an optical tweezers assay, we found that the V8M and Y89D mutations resulted in a significantly reduced force generation (Fig 5). Thus, our results extend the model that nucleotide-dependent conformational changes in the NL are an important mechanical element for force generation by kinesin motors.

Previous work on KIF1A by Nitta et al. (37) examined the effects of mutations in β0, β9 and β10 on KIF1A motor activity and found that while mutations in these three structural elements resulted in relatively normal ATPase activities, mutation of β9 resulted in a decreased velocity in MT-gliding assays. These results support the idea that NL docking, particularly CNB formation, is critical for KIF1A motility, however, these mutations were examined in the context of monomeric motors that contained the catalytic core of KIF1A fused to the NL of the kinesin-1 motor KIF5C. Thus, our work provides the first evidence that NL docking is critical for force generation by KIF1A motors.

### Allostery between force generation and motility properties

KIF1A motors containing V8M or Y89D mutations also exhibited changes in their unloaded motility properties. The speed of both mutant motors was reduced, likely due to allosteric effects of impaired NL docking on motor regions that coordinate and bind nucleotide (S1-PL and S1-S2, Figs 2 and 3). These findings are consistent with previous structural and enzymatic studies suggesting that docking of the NL gates ATPase activity in both kinesin-1 and kinesin-3 motors (30, 37, 49, 50, 68, 69).

The V8M and Y89D mutant motors also exhibited defects in motor-MT interactions as they were less able to engage productively with MTs (lower on-rate) and moved with reduced processivity. These observations appear to conflict with a recent single-molecule characterization of full-length KIF1A (70), which showed that homodimeric human KIF1A motors containing the V8M mutation have an increased landing rate and velocity as compared to the WT motor. However, these apparent discrepancies can be explained by the fact that the V8M mutation relieves auto-inhibition of the full-length motor. That is, the V8M mutation activates the full-length motor in the study of Chiba *et al.* (70) but reduces the MT engagement of our constitutively-active, minimal dimeric motors. Similarly, the V8M mutation increases the velocity of the auto-inhibited full-length motor (70) whereas it reduces the velocity of our minimal dimeric motors (Table 3). Taken together, the data suggest that while the V8M mutation results in a (toxic) gain-of-function in the animal caused by the relief of auto-inhibition (70), the mutant motors are hampered by a reduced MT on-rate, velocity and processivity. Similar effects have been described for other kinesin motors where mutations that relieve autoinhibition can have varying effects on singlemolecule properties *in vitro* and result in gain-of-function phenotypes in cells or animals (70-76).

Finally, for the Y89D mutation, we found that a significant fraction of the mutant motors engaged in non-motile, diffusive MT-binding events and those motors that did undergo processive motility appeared to “wobble” as they walked (Fig 6). These results indicate that the Y89D motor is trapped in an ADP-bound and weakly MT-bound state (16). While the MD simulations predicted that the Y89D mutation would impair ATP hydrolysis but not nucleotide binding (Fig 4C-E), the simulations were carried out on motors strongly bound to MTs in the ATP state. Our results therefore suggest that the Y89D mutation has an additional impact on motility in a stage of the mechanochemical cycle prior to the ATP state, namely, in the motor’s ability to release ADP in response to MT binding.

### Effects on cargo transport and implications for disease

The mechanical and motility properties of KIF1A are likely matched to the cellular functions of this motor and are optimized for transport under physiological conditions. KIF1A motors drive long-range transport of SVPs and DCVs in neurons (4-9) under conditions where teams of 2-4 motors engage with the MT (77, 78). The fast and superprocessive motility of KIF1A motors would be advantageous for long-distance transport and a high-force output may not be required for teams of motors to transport small membrane-bound organelles. The rapid detachment and reattachment of individual motors in response to a hindering load would prevent motors from slowing or stalling and thereby help teams of motors navigate obstacles and ensure fast, continuous transport.

How mutations in KIF1A protein cause disease is still unclear and both loss-of-function and gain-of-function mutations have been linked to human neurodevelopmental and neurodegenerative diseases. V8M and Y89D are *de novo* mutations that manifest in an autosomal dominant manner to cause pure hereditary spastic paraplegia with childhood onset [OMIM #610357, (43)] and Mental retardation, autosomal dominant 9 [OMIM #614255, (44)], respectively. Our results indicate that these mutations result in reduced speed, processivity, landing-rate, and force output of single KIF1A motors and delayed transport driven by teams of mutant motors in an unpolarized cell (Fig 8). Furthermore, our single-molecule motility results suggest that the presence of a mutant motor domain is sufficient to impair the motility properties of heterodimeric WT/mutant motors (Fig 7). It seems likely that in patients, transport driven by these mutant motors is compromised given the long-distances and spatial constraints that characterize transport in neuronal cells.

## MATERIALS AND METHODS

### Structural model and MD simulations of KIF1A-motor complex

Initial coordinates of KIF1A kinesin motor domain in the ATP-bound state (ATP analogue, ANP) and in complex with the tubulin heterodimer were taken from PDB 4UXP (30). The kinesin motor domain sequence was that of *Hs*KIF1A (Uniprot ID Q12756). Missing coordinates, where applicable, were modeled using MODELLER v9.18 (79). A total of 100 models were generated with the following options in MODELLER: variable target function method (VTFM) was set to slow with associated conjugate gradient set to 150 iterations, MD with simulated annealing option was set to slow, and the entire optimization process was repeated twice. The top-scoring model was selected for MD simulations with discrete optimized protein energy (DOPE) score (80) for loop refinement.

Energy minimization and MD simulations were performed with AMBER 18 (University of California San Francisco) and the ff99SB AMBER force field (81). Nucleotide parameters were obtained from (82). Histidine protonation states were assigned based on the their pKa values calculated by PROPKA (83). MD simulations were started from equilibrated structures with at least four independent runs of at least 200 ns each. All simulations were performed in-house on NVIDIA GPU cards with the GPU version of PMEMD (pmemd.cuda). We thank NVIDIA for their gift of GPU card through their Academic GPU seed grant. Trajectory analyses were carried out in R using the Bio3D v2.3-3 package (84).

Residue-residue distance differences between wildtype (WT) and mutant ATP-bound kinesin motor domain in complex with tubulin heterodimer were identified with ensemble difference distance matrix (eDDM) analysis routine (36, 85). For this analysis, a total of 400 conformations were obtained for each state under comparison by extracting 100 equally time-spaced conformations from the last 20 ns of each simulation replicate. Briefly, the eDDM routine reduces the difference between long distances while differences between short distances are kept intact. The significance of residue distance variation between apo and ATP-bound states, and between ATP-bound and mutant states, were evaluated with the Wilcoxon test. Residue pairs showing a p-value <10^-5^ and an average masked distance difference >1 Å were considered statistically significant residue-residue distance differences for further analysis.

### Plasmids

A truncated, constitutively active kinesin-3 [rat KIF1A(1-393)] followed by a leucine zipper was used (16). Point mutations were generated using QuickChange site-directed mutagenesis and all plasmids were verified by DNA sequencing. Motors were tagged with an Avitag for biotinylation and attachment to beads in optical tweezers assays, three tandem monomeric Citrine fluorescent proteins (3xmCit), a monomeric NeonGreen or Halo-FLAGtag for single-molecule imaging assays, or monomeric NeonGreen (mNG)-FRB for inducible cargo dispersion assays in cells (54, 86). The peroxisome-targeting PEX3-mRFP-FKBP construct was a gift from Casper Hoogenraad (Utrecht University (53)).Constructs coding for FRB (DmrA) and FKBP (DmrC) sequences were obtained from ARIAD Pharmaceuticals and are now available from Takara Bio Inc. Plasmids encoding monomeric NeonGreen were obtained from Allele Biotechnology.

A plasmid for *E. coli*-purified KIF1A(393)-LZ was a gift from Dr. Kassandra M. Ori-McKenney (UC Davis (87)). The plasmid was sequenced to ensure that no mutations were present. The construct was amplified by PCR and inserted into pSNAP-tag^®^(T7)-2 vector (New England Biolabs Inc. #N9181S) with a SNAPf-EGFP-6His cassette. Single point mutations in KIF1A were generated by the NEB Q5^®^ site-directed mutagenesis kit (New England Biolabs Inc. #E0554S) and confirmed by sequencing.

### Cell culture, transfection, and lysate preparation

COS-7 (African green monkey kidney fibroblasts, American Type Culture Collection, RRID:CVCL_0224) were grown at 37°C with 5% (vol/vol) CO2 in Dulbecco’s Modified Eagle Medium (Gibco) supplemented with 10% (vol/vol) Fetal Clone III (HyClone) and 2 mM GlutaMAX (L-alanyl-L-glutamine dipeptide in 0.85% NaCl, Gibco). Cells are checked annually for mycoplasma contamination and were authenticated through mass spectrometry (the protein sequences exactly match those in the African green monkey genome). 24 hours after seeding, cells were transfected using TransIT-LT1 transfection reagent (Mirus) and the JF552 HaloTag ligand (Tocris Bioscience) was added to cell culture media to a final concentration of 50 nM. Cells were trypsinized and harvested 24 hours after transfection by low-speed centrifugation at 3000 x g at 4°C for 3 minutes. The pellet was resuspended in cold 1X PBS, centrifuged at 3000 x g at 4°C for 3 minutes, and the pellet was resuspended in 50 μL of cold lysis buffer [25 mM HEPES/KOH, 115 mM potassium acetate, 5 mM sodium acetate, 5 mM MgCl_2_, 0.5 mM EGTA, and 1% (vol/vol) Triton X-100, pH 7.4] with 1 mM ATP, 1 mM phenylmethylsulfonyl fluoride, and 1% (vol/vol) protease inhibitor cocktail (P8340, Sigma-Aldrich). Lysates were clarified by centrifugation at 20,000 x g at 4°C for 10 minutes and lysates were snap frozen in 5 μL aliquots in liquid nitrogen and stored at −80°C.

### Protein Expression and Purification from *E. coli*

Plasmids were transformed into BL21-CodonPlus(DE3)-RIPL competent cells (Agilent Technologies #230280). A single colony was inoculated in 1 mL of terrific broth (TB) with 50 μg/mL carbenicillin and 50 μg/mL chloramphenicol. The 1-mL culture was shaken at 37°C overnight, and then inoculated into 400 mL of TB with 2 μg/mL carbenicillin and 2 μg/mL chloramphenicol. The 400-mL culture was shaken at 37°C for 4-5 hours, and then cooled on ice for 1 hour. IPTG was then added to the culture to final 0.1 mM concentration to induce expression. Afterwards the culture was shaken at 18°C overnight. The cells were harvested by centrifugation at 3000 rcf for 10 minutes at 4°C. The supernatant was discarded, and 5 mL of B-PER™ complete bacterial protein extraction reagent (Thermo Scientific #89821) with 2 mM MgCl_2_, 1 mM EGTA, 1 mM DTT, 0.1 mM ATP, 2 mM PMSF, and 10% glycerol was added to the cell pellet to fully resuspend the cells. The resuspended cells were flash frozen and store at −80°C.

To purify protein, the frozen cells were thawed at 37°C. The solution was nutated at room temperature for 20 minutes and then dounced for 10 strokes on ice to lyse the cells. The cell lysate was cleared by centrifugation at 80,000 rpm for 10 minutes at 4°C using Beckman tabletop centrifuge unit. The lysate was nutated with 200 μL of Ni-NTA resin (Roche cOmplete™ His-Tag purification resin, Millipore Sigma #5893682001) at 4°C for 1 hour. The resin was washed with wash buffer (50 mM HEPES, 300 mM KCl, 2 mM MgCl_2_, 1 mM EGTA, 1 mM DTT, 1 mM PMSF, 0.1 mM ATP, 0.1% Triton X-100, 10% glycerol, pH 7.2), and labeled with 10 μM SNAP-Cell^®^ TMR-Star (New England Biolabs Inc. #S9105S) at room temperature for 10 minutes. The resin was further washed, and the protein was eluted with elution buffer (wash buffer with 250 mM imidazole). The elute was flash frozen and store −80°C.

To remove inactive motors, an MT-binding and -release (MTBR) assay was performed (46). 50 μL of eluted protein was buffer exchanged into low salt buffer (30 mM HEPES, 50 mM KCl, 2 mM MgCl_2_, 1 mM EGTA, 1 mM DTT, and 0.1 mM AMP-PNP) using 0.5-mL Zeba™ spin desalting column (7-kDa MWCO) (Thermo Scientific #89882). AMP-PNP and taxol were added to the flow-through to a final concentration of 1 mM and 10 μM, respectively. After 5 μL of 5 mg/mL taxol-stabilized MTs was added to the mixture, the solution was incubated at room temperature for 5 minutes to allow motors to bind to the MTs. The mixture was then spun through a 100 μl glycerol cushion (PIPES 80 mM, 2 mM MgCl_2_, 1 mM EGTA, 1 mM DTT, 10 μM taxol, and 60% glycerol) by centrifugation at 40000 rpm for 10 minutes at room temperature. Next, the supernatant was removed and the pellet was resuspended in 50 μL high salt buffer (30 mM HEPES, 300 mM KCl, 2 mM MgCl_2_, 1 mM EGTA, 1 mM DTT, 10 μM taxol, 3 mM ATP, and 10% glycerol). The MTs were then removed by centrifugation at 40,000 rpm for 5 minutes at room temperature. Finally, the supernatant was aliquoted and flash frozen in liquid nitrogen and stored at −80°C (MTBR fraction).

### TIRF single-molecule motility assays

MTs were polymerized (tubulin, Cytoskeleton Inc) in BRB80 buffer (80 mM Pipes/KOH pH 6.8, 1 mM MgCl_2_, 1 mM EGTA) supplemented with GTP and MgCl_2_ and incubated for 60 minutes at 37°C. 2 μM taxol in prewarmed BRB80 was added and incubated for 60 minutes to stabilize MTs. MTs were stored in the dark at room temperature for up to 2 weeks. Flow cells were prepared by attaching a #1.5 mm coverslip (Thermo Fisher Scientific) to a glass slide (Thermo Fisher Scientific) using double-sided tape. MTs were diluted in fresh BRB80 buffer supplemented with 10 μM taxol, infused into flow cells, and incubated for four minutes to allow for nonspecific absorption to the glass. Flow cells were then incubated with blocking buffer [0.5 mg/mL casein in imaging buffer supplemented with 10 μM taxol] for four minutes. Flow cells were then infused with motility mixture (0.5–1.0 μL of COS7 cell lysate, 25 μL imaging buffer, 15 μL blocking buffer, 1 mM ATP, 0.5 μL 100 mM DTT, 0.5 μL 20 mg/mL glucose oxidase, 0.5 μL 8 mg/mL catalase, and 0.5 μL 1 M glucose), sealed with molten paraffin wax, and imaged on an inverted Nikon Ti-E/B TIRF microscope with a perfect focus system, a 100x 1.49 NA oil immersion TIRF objective, three 20 mW diode lasers (488 nm, 561 nm, and 640 nm) and an EMCCD camera (iXon+ DU879; Andor). Image acquisition was controlled using Nikon software and all assays were performed at room temperature. To optimize the single-molecule imaging conditions for KIF1A motors in COS-7 cell lysates, the following imaging buffers were utilized: P12 (12 mM Pipes/KOH pH 6.8, 1 mM MgCl_2_, 1 mM EGTA), BRB40 (40 mM Pipes/KOH pH 6.8, 1 mM MgCl_2_, 1 mM EGTA), BRB80 (80 mM Pipes/KOH pH 6.8, 1 mM MgCl_2_, 1 mM EGTA), or PERM (25mM HEPES/KOH, 115mM potassium acetate, 5mM sodium acetate, 5mM MgCl_2_, and 0.5mM EGTA, pH 7.4). Motility assays were carried out in BRB40 buffer with MTs 35-75 μm in length.

Motility data were analyzed by first generating maximum intensity projections to identify MT tracks (width = 3 pixels) and then generating kymographs in ImageJ (National Institutes of Health). All motility events that lasted for at least three frames were analyzed. As many of these events end when the motor reaches the end of a MT, the reported run lengths are an underestimation of the motor’s processivity. The reported run lengths are also limited by the length of the MTs in the imaging chamber. For each motor construct, the velocities and run lengths were binned and a histogram was generated by plotting the number of motility events for each bin. At least 150 motility events were quantified for each motor across three independent trials and summarized as a histogram or dot plot. A corresponding Gaussian or exponential distribution was overlaid on each histogram plot using rate and shape parameters derived from fitting the cumulative distributions. Motor velocities were fit to a Gaussian cumulative distribution as previously described (41) and a one-way analysis of variance test was used to assess whether velocity distributions were significantly different between motors. Motor run lengths were fit to an exponential decay cumulative distribution where appropriate as previously described (41) and a Kruskal-Wallis one-way analysis of variance was used to assess whether run length distributions were significantly different between motors. For run lengths histograms that did not follow an exponential distribution, median values with percentiles (25%, 75%) were calculated.

Single-molecule TIRF motility studies of *E. coli* expressed and purified KIF1A motors was performed as described (46). Motility assays were carried out in BRB40 buffer with MTs 10-35 μm in length. For each movie, a total of 600 frames was acquired with and acquisition time of 200 ms per frame.

### Optical tweezers assay

The polystyrene trapping beads, MTs and slides were prepared as described previously (46). Briefly, polystyrene beads with an average diameter of 500 nm (Bangs Laboratories Inc. #PC02002) were coated with streptavidin and α-casein, or with an anti-GFP antibody and α-casein. Coverslips (Zeiss #474030-9000-000) were cleaned with 25% HNO3 and 2 M NaOH, washed with ddH2O, air dried, and stored at 4°C. The flow chamber was assembled with a glass slide, parafilm stripes, and a cleaned coverslip as described (46). MTs with incorporated biotinylated tubulin were attached to the cover glass surface via α-casein-biotin and streptavidin. Control cell lysate without KIF1A expression was tested to ensure there were no non-specific interactions between other endogenous motors in the lysate and the beads. 100 beads were tested and no force generation was observed under the same experimental conditions used for cell lysates containing tagged KIF1A constructs. Cell lysate with KIF1A was pre-diluted 50-200x, while the MTBR fraction of *E. coli*-expressed KIF1A was pre-diluted 200-5000x. 1 μL of the predilution was incubated with 0.4 μL beads on ice for 15 minutes. For experiments with the cell lysate, the lysate was pre-diluted so that less than 10% of the beads showed force generation; for experiments with the *E. coli*-expressed KIF1A, the solution was diluted so that less than 30% of the beads tested showed force generation events. Finally, the protein-bead mixture was diluted in 40 μL assay buffer (60 mM HEPES, 50 mM KoAc, 2 mM MgCl_2_, 1 mM EGTA, 1 mM DTT, 10 μM taxol, 2 mM ATP, 50 mM glucose, gloxy, 1.25 mg/ml α-casein, 10% glycerol) and flowed into the slide chamber. All optical trapping experiments were performed with a LUMICKS C-Trap^®^, which combines optical tweezers with 3-color TIRF microscopy and interference reflection microscopy (IRM) to visualize unlabeled MTs (88).

### Inducible peroxisome dispersion assay

Plasmids for expression of WT or mutant rat KIF1A(339)-LZ motors tagged with monomeric NeonGreen and an FRB domain were cotransfected into COS-7 cells with a plasmid for expression of PEX3-mRFP-FKBP at a ratio of 6:1 with TransITLT1 transfection reagent (Mirus). Eight hours after transfection, rapamycin (Calbiochem, Millipore Sigma) was added to final concentration of 44 nM to promote FRB and FKBP heterodimerization and recruitment of motors to peroxisomes. 0, 5, 10, or 30 minutes after addition of rapamycin and recruitment of motors to the surface of peroxisomes, cells were fixed with 3.7% formaldehyde (Thermo Fisher Scientific) in 1X PBS for 10 minutes, quenched in 50 mM ammonium chloride in PBS for 5 minutes, and permeabilized in 0.2% Triton X-100 in PBS for 5 minutes. Coverslips were mounted in ProlongGold (Invitrogen) and imaged using an inverted epifluorescence microscope (Nikon TE2000E) with a 40x/0.75 NA objective and a CoolSnapHQ camera (Photometrics). Only cells expressing low levels of motor-mNG-FRB were imaged and included in quantification. The phenotype of cargo dispersion was scored as clustered, partial dispersion, diffuse dispersion, or peripheral dispersion based on the signal localization in the PEX3 (peroxisome) channel. The data for each construct across three independent trials is summarized as a stacked bar plot.

## ACKNOWLEDGEMENTS

BGB was supported by the NIH Cellular and Molecular Biology Training Grant T32-GM007315, a Graduate Student Research Fellowship from NSF DGE 1256260, and a Rackham Predoctoral Fellowship from the University of Michigan. SJ was supported by the Qatar Research Leadership Program. Work in the laboratory of KJV is supported by NIH grants R01GM070862 and R35GM131744. DS is supported by NIH grant R01GM136822. LR and AG are supported by the NIH grants R01GM098469 and R01NS114636.

## AUTHOR CONTRIBUTIONS

BGB, SJ, LR, DS, KJV, and AG designed the research; BGB, SJ and LR performed the research; BGB produced WT and mutant KIF1A motors expressed in COS-7 cells and LR produced and purified *E. coli*-expressed proteins. BGB and LR analyzed experimental data and SJ and DS performed MD simulations. BGB, SJ, LR, DS, KJV, and AG wrote the manuscript.

## SUPPLEMENTAL FIGURES

**Figure S1.**
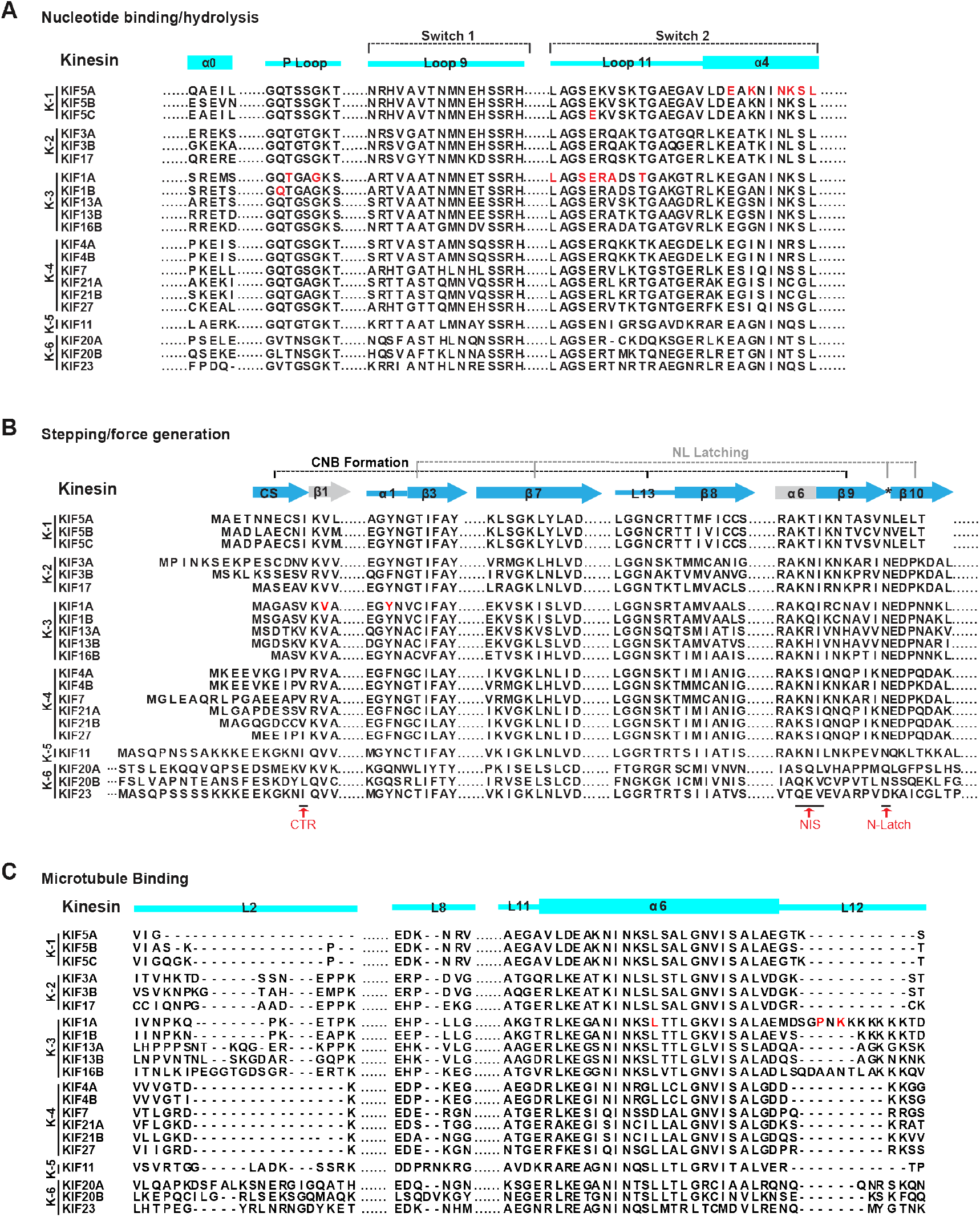
Sequence alignment of functional elements of the kinesin motor domain across the kinesin superfamily. (A-C) Alignment of the motor domain sequences from the indicated human members of the kinesin-1, −2, −3, −4, −5, −6, and −10 families. Secondary structural elements important for (A) MT binding, (B) nucleotide binding/hydrolysis, and (C) force generation/stepping are illustrated. Red text denotes identified mutations associated with neurodevelopmental and/or neurodegenerative disorders. CTR, C-terminal residue of the CS. NIS, NL initiation sequence. N-Latch, asparagine (N) latch.

**Figure S2.**
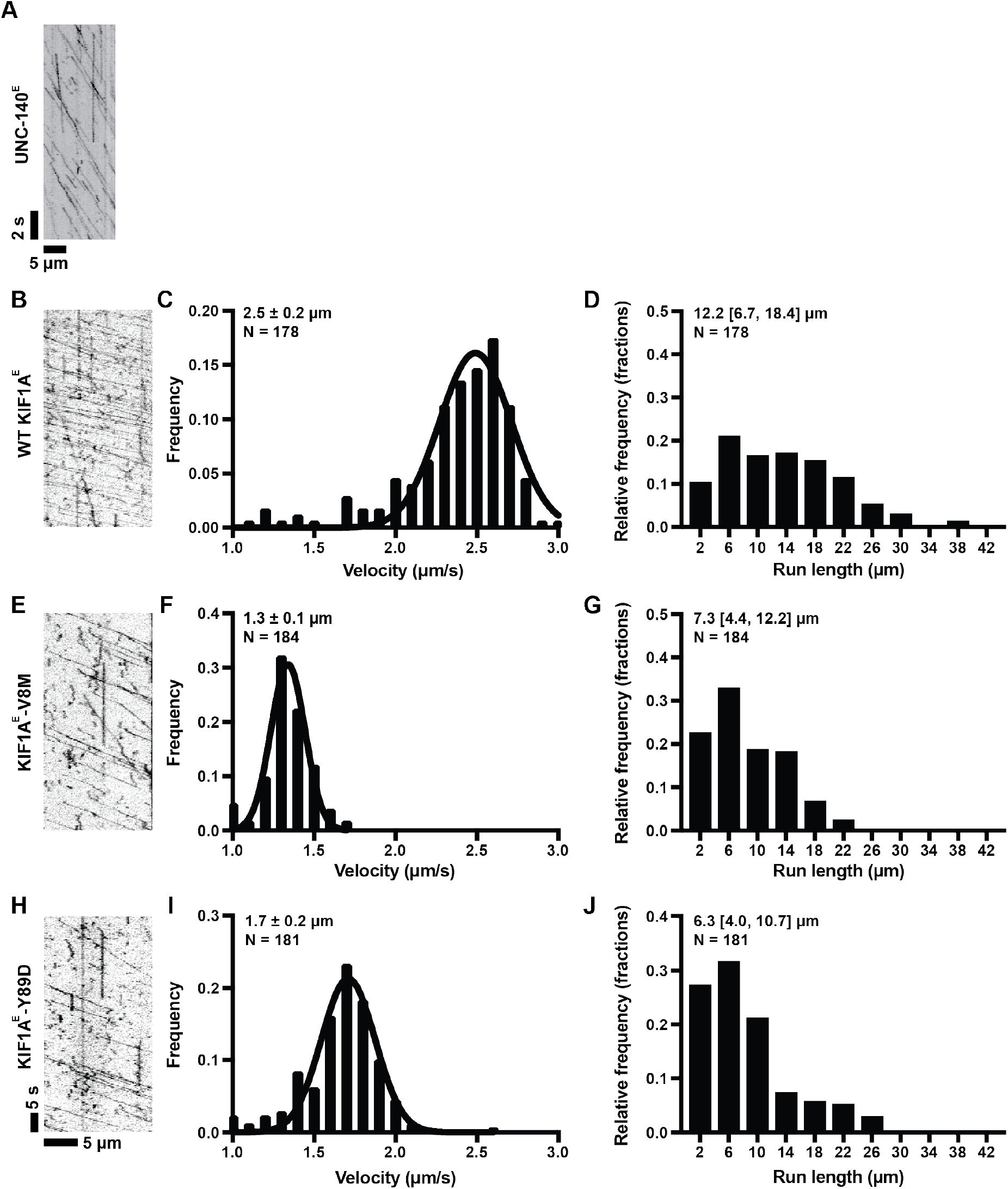
Velocity and processivity of *E. coli*-expressed WT and mutant KIF1A. (A) Example kymograph of UNC-104^E^ purified from *E. coli* bacteria. (B-D) Example kymograph of WT KIF1A^E^. From kymographs, single-motor velocities (C) and run lengths (D) were determined. The mean values ± SEM (for velocities) and the median with quartiles (for run lengths) are indicated on each graph. (E-G) As in *C-D* but for the KIF1A^E^-V8M mutant. (H-J) As in *C-D* but for the KIF1A^E^-Y89D mutant.

**Figure S3.**
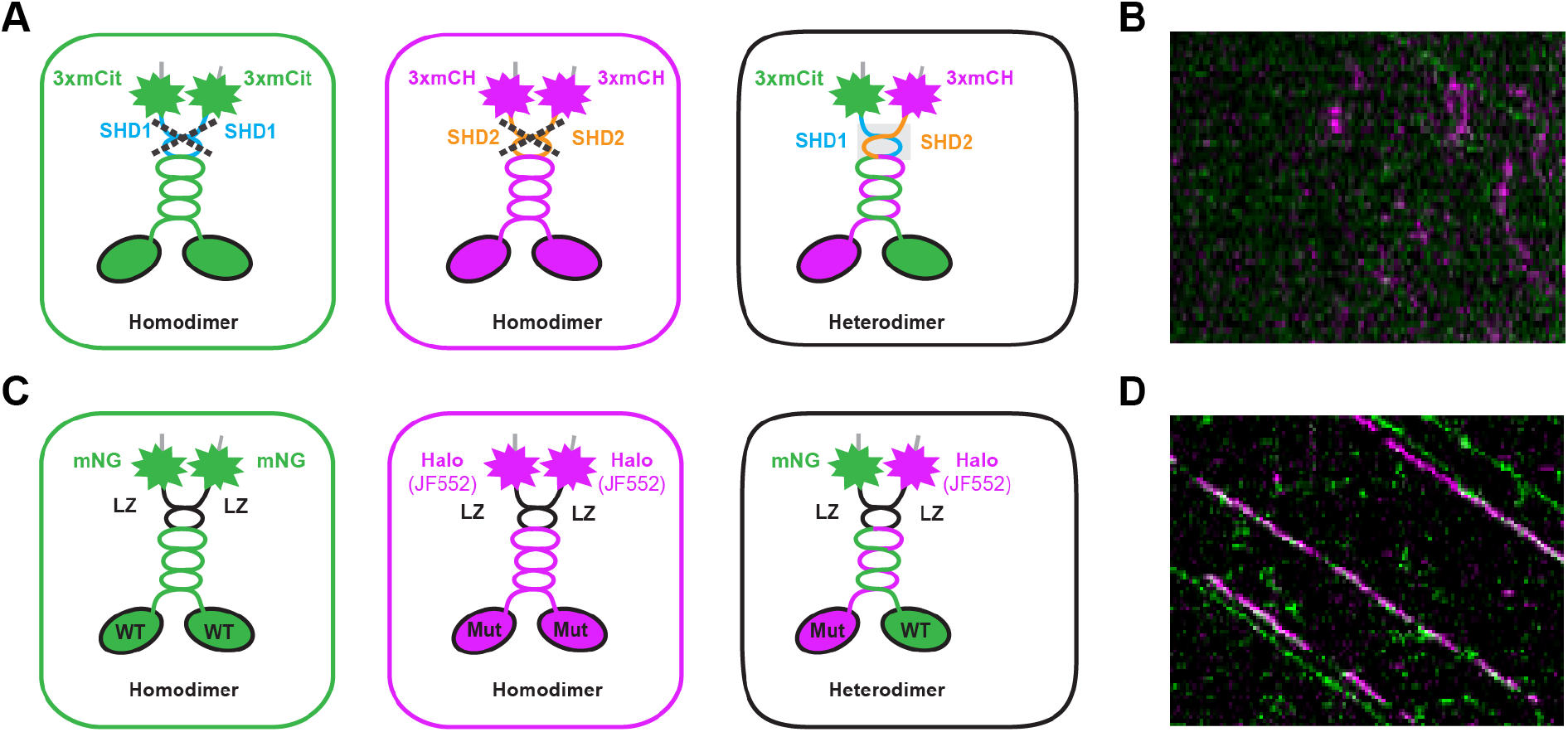
Strategies for designing heterodimeric motors. (A,B) Formation of heterodimeric motors using a synthetic heterodimerization (SHD) sequence. (A) SHD (Albracht et al., 2014 JBC) was fused to the C-terminus of KIF1A(1-393). Coiled-coil prediction software was used to ensure that the SHD sequences were placed in register with the native KIF1A heptad repeat (Marcoil). One KIF1A(1-393)-SHD sequence was fused to three tandem monomeric citrine florescent proteins [KIF1A(393)-SHD1-3xmCit] and the other was fused to three tandem monomeric Cherry proteins [KIF1A(393)-SHD2-3xmCH]. Unlike the leucine zipper sequence of GCN4, SHD1 and SHD2 sequences are not expected to homodimerize (left and middle) and instead are expected to form a heterodimer (right). To test this, lysates of COS-7 cells co-transfected with plasmids coding for KIF1A(393)-SHD1-3xmCit and KIF1A(393)-SHD2-3xmCH motors were subjected to single-molecule imaging using TIRF microscopy. (B) Representative kymograph of a TIRF single-molecule assay of lysates from COS-7 cells cotransfected with KIF1A(1-393)-SHD1-3xmCit and KIF1A(1-393)-SHD2-3xmCH plasmids. Time is displayed on the y-axis (bar, 4 s) and distance displayed on the x-axis (bar, 4 μm). Very few heterodimeric (magenta/green) spots were detected. The few heterodimeric events observed were short and non-processive, unlike the fast, super-processive motility of stable dimeric KIF1A motors. (C,D) Formation of heterodimeric motors using a leucine zipper (LZ) sequence. (C) The LZ sequence of GCN4 was fused to the C-terminus of KIF1A(1-393) (Hammond 2009 PLoS Biol). Coiled-coil prediction software was used to ensure that the LZ sequences were placed in register with the native KIF1A heptad repeat (Marcoil). Motors were fused to either monomeric NeonGreen (mNG) or Halo-FLAG fluorescently tagged with JF552. Three populations of motors are expected in TIRF single-molecule assays: homodimeric Halo-FLAG motors, homodimeric mNG motors, and heterodimeric Halo-FLAG/mNG motors. (D) Representative kymograph of TIRF single-molecule assays of lysates from COS-7 cells cotransfected with KIF1A(1-393)-LZ-mNG and KIF1A(1-393)-LZ-Halo-FLAG. Time is displayed on the x-axis (bar, 4 s) and distance displayed on the y-axis (bar, 4 μm). Heterodimeric motors (green/magenta) showed fast, superprocessive runs typical of KIF1A motors.

**Figure S4.**
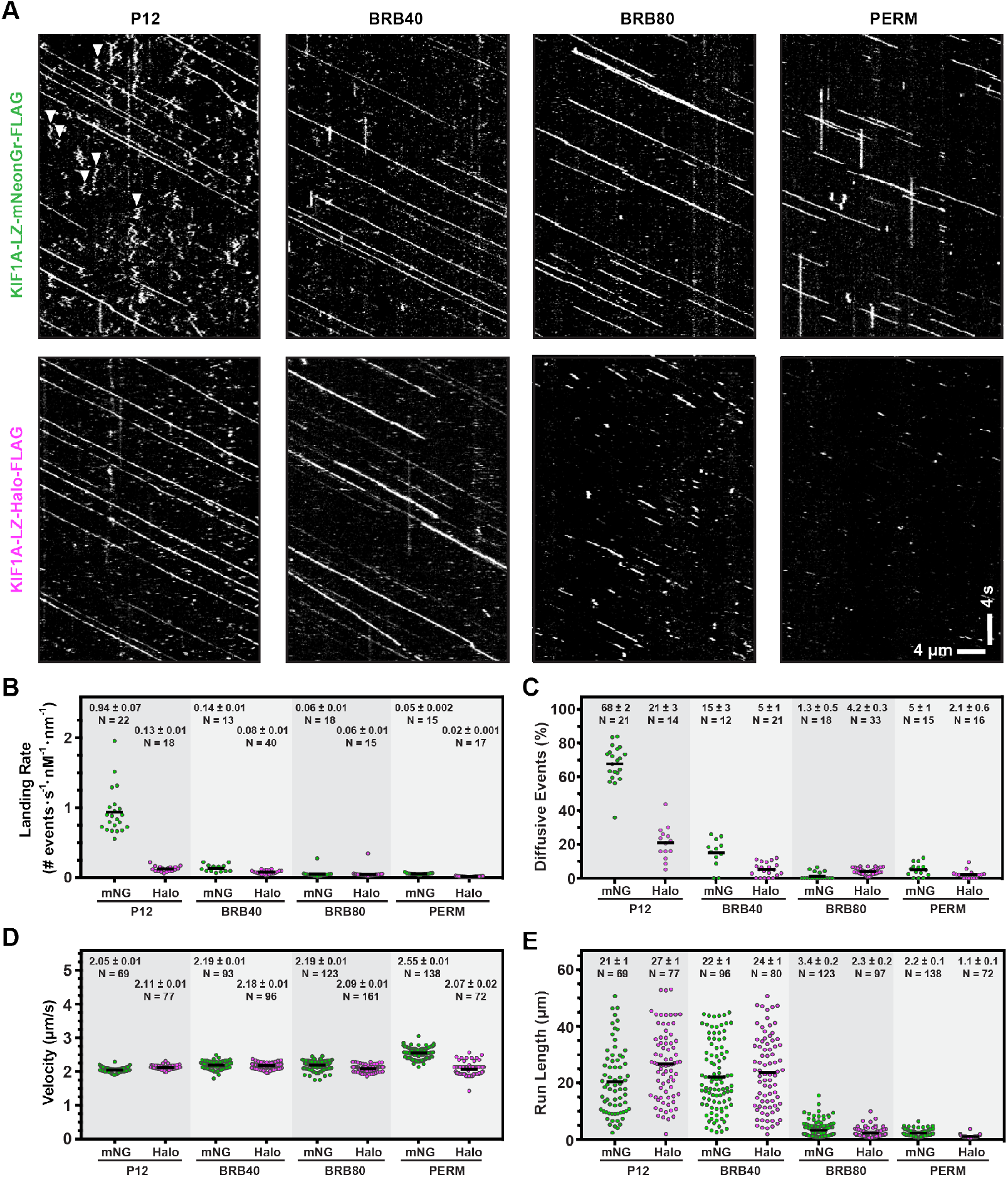
Influence of fluorescence tag and buffer conditions on KIF1A motility. (A) Motility properties of KIF1A motors dimerized by a leucine zipper sequence (LZ) and fused at their C-termini to monomeric NeonGreen (mNG) or Halo-FLAG/JF552. Motors in COS-7 lysates were analyzed in standard single-molecule motility assays using TIRF microscopy. Representative kymographs are shown with time displayed on the y-axis (bar, 4 s) and distance displayed on the x-axis (bar, 4 μm). White arrowheads indicate motility events scored as a diffusive. (B-E) Quantification of motility properties. From the kymographs, single-motor landing rates (B) [motility events (diffusive and processive) with dwell times longer than 400 ms], frequency of diffusive events (C) (net displacement <300 nm, dwell time >400ms, velocity (D), and run length (E) were determined and the data for each population is plotted as a dot plot. Each dot represents a single motor. Consistent with previous studies (Norris et al., 2015), buffer conditions had little effect on velocity but did affect the other parameters in a tag-dependent manner. Further experiments were carried out in BRB40 buffer.

## REFERENCES

1. Hirokawa N, Noda Y, Tanaka Y, Niwa S. Kinesin superfamily motor proteins and intracellular transport. Nat Rev Mol Cell Biol. 2009;10(10):682–96.

2. Siddiqui N, Straube A. Intracellular Cargo Transport by Kinesin-3 Motors. Biochemistry (Mosc). 2017;82(7):803–15.

3. Gabrych DR, Lau VZ, Niwa S, Silverman MA. Going Too Far Is the Same as Falling Short(dagger): Kinesin-3 Family Members in Hereditary Spastic Paraplegia. Front Cell Neurosci. 2019;13:419.

4. Barkus RV, Klyachko O, Horiuchi D, Dickson BJ, Saxton WM. Identification of an axonal kinesin-3 motor for fast anterograde vesicle transport that facilitates retrograde transport of neuropeptides. Mol Biol Cell. 2008;19(1):274–83.

5. Hall DH, Hedgecock EM. Kinesin-related gene unc-104 is required for axonal transport of synaptic vesicles in C. elegans. Cell. 1991;65(5):837–47.

6. Lo KY, Kuzmin A, Unger SM, Petersen JD, Silverman MA. KIF1A is the primary anterograde motor protein required for the axonal transport of dense-core vesicles in cultured hippocampal neurons. Neurosci Lett. 2011;491(3):168–73.

7. Okada Y, Yamazaki H, Sekine-Aizawa Y, Hirokawa N. The neuron-specific kinesin superfamily protein KIF1A is a unique monomeric motor for anterograde axonal transport of synaptic vesicle precursors. Cell. 1995;81(5):769–80.

8. Yonekawa Y, Harada A, Okada Y, Funakoshi T, Kanai Y, Takei Y, et al. Defect in synaptic vesicle precursor transport and neuronal cell death in KIF1A motor protein-deficient mice. J Cell Biol. 1998;141(2):431–41.

9. Zahn TR, Angleson JK, MacMorris MA, Domke E, Hutton JF, Schwartz C, et al. Dense core vesicle dynamics in Caenorhabditis elegans neurons and the role of kinesin UNC-104. Traffic. 2004;5(7):544–59.

10. Eggermann K, Gess B, Hausler M, Weis J, Hahn A, Kurth I. Hereditary Neuropathies. Dtsch Arztebl Int. 2018;115(6):91–7.

11. Guo Y, Chen Y, Yang M, Xu X, Lin Z, Ma J, et al. A Rare KIF1A Missense Mutation Enhances Synaptic Function and Increases Seizure Activity. Front Genet. 2020;11:61.

12. Martin PB, Hicks AN, Holbrook SE, Cox GA. Overlapping spectrums: The clinicogenetic commonalities between Charcot-Marie-Tooth and other neurodegenerative diseases. Brain Res. 2020;1727:146532.

13. Nemani T, Steel D, Kaliakatsos M, DeVile C, Ververi A, Scott R, et al. KIF1A-related disorders in children: A wide spectrum of central and peripheral nervous system involvement. J Peripher Nerv Syst. 2020;.

14. Samanta D, Gokden M. PEHO syndrome: KIF1A mutation and decreased activity of mitochondrial respiratory chain complex. J Clin Neurosci. 2019;61:298–301.

15. Wang J, Zhang Q, Chen Y, Yu S, Wu X, Bao X. Rett and Rett-like syndrome: Expanding the genetic spectrum to KIF1A and GRIN1 gene. Mol Genet Genomic Med. 2019;7(11):e968.

16. Hammond JW, Cai D, Blasius TL, Li Z, Jiang Y, Jih GT, et al. Mammalian Kinesin-3 motors are dimeric in vivo and move by processive motility upon release of autoinhibition. PLoS Biol. 2009;7(3):e72.

17. Scarabelli G, Soppina V, Yao XQ, Atherton J, Moores CA, Verhey KJ, et al. Mapping the Processivity Determinants of the Kinesin-3 Motor Domain. Biophys J. 2015;109(8):1537–40.

18. Soppina V, Norris SR, Dizaji AS, Kortus M, Veatch S, Peckham M, et al. Dimerization of mammalian kinesin-3 motors results in superprocessive motion. Proc Natl Acad Sci U S A. 2014;111(15):5562–7.

19. Svoboda K, Block SM. Force and velocity measured for single kinesin molecules. Cell. 1994;77(5):773–84.

20. Asbury CL, Fehr AN, Block SM. Kinesin moves by an asymmetric hand-over-hand mechanism. Science. 2003;302(5653):2130–4.

21. Guydosh NR, Block SM. Direct observation of the binding state of the kinesin head to the microtubule. Nature. 2009;461(7260):125–8.

22. Hwang W, Karplus M. Structural basis for power stroke vs. Brownian ratchet mechanisms of motor proteins. Proc Natl Acad Sci U S A. 2019;116(40):19777–85.

23. Ramaiya A, Roy B, Bugiel M, Schaffer E. Kinesin rotates unidirectionally and generates torque while walking on microtubules. Proc Natl Acad Sci U S A. 2017;114(41):10894–9.

24. Brenner S, Berger F, Rao L, Nicholas MP, Gennerich A. Force production of human cytoplasmic dynein is limited by its processivity. Sci Adv. 2020;6(15):eaaz4295.

25. Case RB, Rice S, Hart CL, Ly B, Vale RD. Role of the kinesin neck linker and catalytic core in microtubule-based motility. Curr Biol. 2000;10(3):157–60.

26. Hwang W, Lang MJ, Karplus M. Kinesin motility is driven by subdomain dynamics. Elife. 2017;6.

27. Khalil AS, Appleyard DC, Labno AK, Georges A, Karplus M, Belcher AM, et al. Kinesin’s cover neck bundle folds forward to generate force. Proc Natl Acad Sci USA. 2008;105(49):19247–52.

28. Rice S, Lin AW, Safer D, Hart CL, Naber N, Carragher BO, et al. A structural change in the kinesin motor protein that drives motility. Nature. 1999;402(6763):778–84.

29. Hwang W, Lang MJ, Karplus M. Force generation in kinesin hinges on cover-neck bundle formation. Structure. 2008;16(1):62–71.

30. Atherton J, Farabella I, Yu IM, Rosenfeld SS, Houdusse A, Topf M, et al. Conserved mechanisms of microtubule-stimulated ADP release, ATP binding, and force generation in transport kinesins. Elife. 2014;3:e03680.

31. Atherton J, Yu IM, Cook A, Muretta JM, Joseph A, Major J, et al. The divergent mitotic kinesin MKLP2 exhibits atypical structure and mechanochemistry. Elife. 2017;6.

32. Goulet A, Behnke-Parks WM, Sindelar CV, Major J, Rosenfeld SS, Moores CA. The structural basis of force generation by the mitotic motor kinesin-5. J Biol Chem. 2012;287(53):44654–66.

33. Goulet A, Major J, Jun Y, Gross SP, Rosenfeld SS, Moores CA. Comprehensive structural model of the mechanochemical cycle of a mitotic motor highlights molecular adaptations in the kinesin family. Proc Natl Acad Sci U S A. 2014;111(5):1837–42.

34. Hesse WR, Steiner M, Wohlever ML, Kamm RD, Hwang W, Lang MJ. Modular aspects of kinesin force generation machinery. Biophys J. 2013;104(9):1969–78.

35. von Loeffelholz O, Moores CA. Cryo-EM structure of the Ustilago maydis kinesin-5 motor domain bound to microtubules. J Struct Biol. 2019;207(3):312–6.

36. Budaitis BG, Jariwala S, Reinemann DN, Schimert KI, Scarabelli G, Grant BJ, et al. Neck linker docking is critical for Kinesin-1 force generation in cells but at a cost to motor speed and processivity. Elife. 2019;8.

37. Nitta R, Okada Y, Hirokawa N. Structural model for strain-dependent microtubule activation of Mg-ADP release from kinesin. Nat Struct Mol Biol. 2008;15(10):1067–75.

38. Ren J, Zhang Y, Wang S, Huo L, Lou J, Feng W. Structural Delineation of the Neck Linker of Kinesin-3 for Processive Movement. J Mol Biol. 2018;430(14):2030–41.

39. Arpag G, Norris SR, Mousavi SI, Soppina V, Verhey KJ, Hancock WO, et al. Motor Dynamics Underlying Cargo Transport by Pairs of Kinesin-1 and Kinesin-3 Motors. Biophys J. 2019;116(6):1115–26.

40. Arpag G, Shastry S, Hancock WO, Tuzel E. Transport by populations of fast and slow kinesins uncovers novel family-dependent motor characteristics important for in vivo function. Biophys J. 2014;107(8):1896–904.

41. Norris SR, Soppina V, Dizaji AS, Schimert KI, Sept D, Cai D, et al. A method for multiprotein assembly in cells reveals independent action of kinesins in complex. J Cell Biol. 2014;207(3):393–406.

42. Tomishige M, Klopfenstein DR, Vale RD. Conversion of Unc104/KIF1A kinesin into a processive motor after dimerization. Science. 2002;297(5590):2263–7.

43. Iqbal Z, Rydning SL, Wedding IM, Koht J, Pihlstrom L, Rengmark AH, et al. Targeted high throughput sequencing in hereditary ataxia and spastic paraplegia. PLoS One. 2017;12(3):e0174667.

44. ClinVar; [VCV000224157.1] [Internet]. [cited April 8, 2020]. Available from: https://www.ncbi.nlm.nih.gov/clinvar/variation/VCV000224157.1

45. Richard J, Kim ED, Nguyen H, Kim CD, Kim S. Allostery Wiring Map for Kinesin Energy Transduction and Its Evolution. J Biol Chem. 2016;291(40):20932–45.

46. Rao L, Berger F, Nicholas MP, Gennerich A. Molecular mechanism of cytoplasmic dynein tension sensing. Nat Commun. 2019;10(1):3332.

47. Nicholas MP, Hook P, Brenner S, Wynne CL, Vallee RB, Gennerich A. Control of cytoplasmic dynein force production and processivity by its C-terminal domain. Nat Commun. 2015;6:6206.

48. Soppina V, Verhey KJ. The family-specific K-loop influences the microtubule on-rate but not the superprocessivity of kinesin-3 motors. Mol Biol Cell. 2014;25(14):2161–70.

49. Cao L, Wang W, Jiang Q, Wang C, Knossow M, Gigant B. The structure of apo-kinesin bound to tubulin links the nucleotide cycle to movement. Nat Commun. 2014;5:5364.

50. Shang Z, Zhou K, Xu C, Csencsits R, Cochran JC, Sindelar CV. High-resolution structures of kinesin on microtubules provide a basis for nucleotide-gated force-generation. Elife. 2014;3:e04686.

51. Kikkawa M, Sablin EP, Okada Y, Yajima H, Fletterick RJ, Hirokawa N. Switch-based mechanism of kinesin motors. Nature. 2001;411(6836):439–45.

52. Rao L, Romes EM, Nicholas MP, Brenner S, Tripathy A, Gennerich A, et al. The yeast dynein Dyn2-Pac11 complex is a dynein dimerization/processivity factor: structural and single-molecule characterization. Mol Biol Cell. 2013;24(15):2362–77.

53. Kapitein LC, Schlager MA, van der Zwan WA, Wulf PS, Keijzer N, Hoogenraad CC. Probing intracellular motor protein activity using an inducible cargo trafficking assay. Biophys J. 2010;99(7):2143–52.

54. Schimert KI, Budaitis BG, Reinemann DN, Lang MJ, Verhey KJ. Intracellular cargo transport by single-headed kinesin motors. Proc Natl Acad Sci U S A. 2019;116(13):6152–61.

55. Efremov AK, Radhakrishnan A, Tsao DS, Bookwalter CS, Trybus KM, Diehl MR. Delineating cooperative responses of processive motors in living cells. Proc Natl Acad Sci U S A. 2014;111(3):E334–43.

56. Lessard DV, Zinder OJ, Hotta T, Verhey KJ, Ohi R, Berger CL. Polyglutamylation of tubulin’s C-terminal tail controls pausing and motility of kinesin-3 family member KIF1A. J Biol Chem. 2019;294(16):6353–63.

57. Rogers KR, Weiss S, Crevel I, Brophy PJ, Geeves M, Cross R. KIF1D is a fast non-processive kinesin that demonstrates novel K-loop-dependent mechanochemistry. EMBO J. 2001;20(18):5101–13.

58. Siddiqui N, Zwetsloot AJ, Bachmann A, Roth D, Hussain H, Brandt J, et al. PTPN21 and Hook3 relieve KIF1C autoinhibition and activate intracellular transport. Nat Commun. 2019;10(1):2693.

59. Carter NJ, Cross RA. Mechanics of the kinesin step. Nature. 2005;435(7040):308–12.

60. Andreasson JO, Shastry S, Hancock WO, Block SM. The Mechanochemical Cycle of Mammalian Kinesin-2 KIF3A/B under Load. Curr Biol. 2015;25(9):1166–75.

61. Milic B, Andreasson JOL, Hogan DW, Block SM. Intraflagellar transport velocity is governed by the number of active KIF17 and KIF3AB motors and their motility properties under load. Proc Natl Acad Sci U S A. 2017;114(33):E6830–E8.

62. Schroeder HW, 3rd, Hendricks AG, Ikeda K, Shuman H, Rodionov V, Ikebe M, et al. Force-dependent detachment of kinesin-2 biases track switching at cytoskeletal filament intersections. Biophys J. 2012;103(1):48–58.

63. Korneev MJ, Lakamper S, Schmidt CF. Load-dependent release limits the processive stepping of the tetrameric Eg5 motor. Eur Biophys J. 2007;36(6):675–81.

64. Shimamoto Y, Forth S, Kapoor TM. Measuring Pushing and Braking Forces Generated by Ensembles of Kinesin-5 Crosslinking Two Microtubules. Dev Cell. 2015;34(6):669–81.

65. Valentine MT, Block SM. Force and premature binding of ADP can regulate the processivity of individual Eg5 dimers. Biophys J. 2009;97(6):1671–7.

66. Valentine MT, Fordyce PM, Krzysiak TC, Gilbert SP, Block SM. Individual dimers of the mitotic kinesin motor Eg5 step processively and support substantial loads in vitro. Nat Cell Biol. 2006;8(5):470–6.

67. Ren J, Wang S, Chen H, Wang W, Huo L, Feng W. Coiled-coil 1-mediated fastening of the neck and motor domains for kinesin-3 autoinhibition. Proc Natl Acad Sci US A. 2018;115(51):E11933–E42.

68. Hahlen K, Ebbing B, Reinders J, Mergler J, Sickmann A, Woehlke G. Feedback of the kinesin-1 neck-linker position on the catalytic site. J Biol Chem. 2006;281(27):18868–77.

69. Muretta JM, Jun Y, Gross SP, Major J, Thomas DD, Rosenfeld SS. The structural kinetics of switch-1 and the neck linker explain the functions of kinesin-1 and Eg5. Proc Natl Acad Sci U S A. 2015;112(48):E6606–13.

70. Chiba K, Takahashi H, Chen M, Obinata H, Arai S, Hashimoto K, et al. Disease-associated mutations hyperactivate KIF1A motility and anterograde axonal transport of synaptic vesicle precursors. Proc Natl Acad Sci U S A. 2019;116(37):18429–34.

71. Cheng L, Desai J, Miranda CJ, Duncan JS, Qiu W, Nugent AA, et al. Human CFEOM1 mutations attenuate KIF21A autoinhibition and cause oculomotor axon stalling. Neuron. 2014;82(2):334–49.

72. Imanishi M, Endres NF, Gennerich A, Vale RD. Autoinhibition regulates the motility of the C. elegans intraflagellar transport motor OSM-3. J Cell Biol. 2006;174(7):931–7.

73. Moua P, Fullerton D, Serbus LR, Warrior R, Saxton WM. Kinesin-1 tail autoregulation and microtubule-binding regions function in saltatory transport but not ooplasmic streaming. Development. 2011;138(6):1087–92.

74. van der Vaart B, van Riel WE, Doodhi H, Kevenaar JT, Katrukha EA, Gumy L, et al. CFEOM1-associated kinesin KIF21A is a cortical microtubule growth inhibitor. Dev Cell. 2013;27(2):145–60.

75. Kelliher MT, Yue Y, Ng A, Kamiyama D, Huang B, Verhey KJ, et al. Autoinhibition of kinesin-1 is essential to the dendrite-specific localization of Golgi outposts. J Cell Biol. 2018;217(7):2531–47.

76. Niwa S, Lipton DM, Morikawa M, Zhao C, Hirokawa N, Lu H, et al. Autoinhibition of a Neuronal Kinesin UNC-104/KIF1A Regulates the Size and Density of Synapses. Cell Rep. 2016;16(8):2129–41.

77. Hayashi K, Hasegawa S, Sagawa T, Tasaki S, Niwa S. Non-invasive force measurement reveals the number of active kinesins on a synaptic vesicle precursor in axonal transport regulated by ARL-8. Phys Chem Chem Phys. 2018;20(5):3403–10.

78. Hayashi K, Tsuchizawa Y, Iwaki M, Okada Y. Application of the fluctuation theorem for non-invasive force measurement in living neuronal axons. Mol Biol Cell. 2018:mbcE18010022.

79. Sali A, Blundell TL. Comparative protein modelling by satisfaction of spatial restraints. J Mol Biol. 1993;234(3):779–815.

80. Shen MY, Sali A. Statistical potential for assessment and prediction of protein structures. Protein Sci. 2006;15(11):2507–24.

81. Hornak V, Abel R, Okur A, Strockbine B, Roitberg A, Simmerling C. Comparison of multiple Amber force fields and development of improved protein backbone parameters. Proteins. 2006;65(3):712–25.

82. Meagher KL, Redman LT, Carlson HA. Development of polyphosphate parameters for use with the AMBER force field. J Comput Chem. 2003;24(9):1016–25.

83. Li H, Robertson AD, Jensen JH. Very fast empirical prediction and rationalization of protein pKa values. Proteins. 2005;61(4):704–21.

84. Skjaerven L, Yao XQ, Scarabelli G, Grant BJ. Integrating protein structural dynamics and evolutionary analysis with Bio3D. BMC Bioinformatics. 2014;15:399.

85. Muretta JM, Reddy BJN, Scarabelli G, Thompson AF, Jariwala S, Major J, et al. A posttranslational modification of the mitotic kinesin Eg5 that enhances its mechanochemical coupling and alters its mitotic function. Proc Natl Acad Sci U S A. 2018;115(8):E1779–E88.

86. Engelke MF, Winding M, Yue Y, Shastry S, Teloni F, Reddy S, et al. Engineered kinesin motor proteins amenable to small-molecule inhibition. Nat Commun. 2016;7:11159.

87. Monroy BY, Sawyer DL, Ackermann BE, Borden MM, Tan TC, Ori-McKenney KM. Competition between microtubule-associated proteins directs motor transport. Nat Commun. 2018;9(1):1487.

88. Mahamdeh M, Simmert S, Luchniak A, Schaffer E, Howard J. Label-free high-speed wide-field imaging of single microtubules using interference reflection microscopy. J Microsc. 2018;272(1):60–6.

